# Pathogen genetic control of transcriptome variation in the *Arabidopsis thaliana* – *Botrytis cinerea* pathosystem

**DOI:** 10.1101/577585

**Authors:** Nicole E. Soltis, Wei Zhang, Jason A. Corwin, Susanna Atwell, Daniel J. Kliebenstein

## Abstract

Disease symptoms arise from the interaction of the host and pathogen genomes. However, little is known about how genetic variation in the interaction modulates both organisms’ transcriptomes, especially in complex interactions like those between generalist pathogens and their plant hosts. To begin mapping how polygenic pathogen variation influences both organisms’ transcriptomes, we used the *Botrytis cinerea* - *Arabidopsis thaliana* pathosystem. We measured the co-transcriptome across a genetically diverse collection of 96 *B. cinerea* isolates infected on the Arabidopsis wildtype, Col-0. Using the *B. cinerea* genomic variation, we performed genome-wide association (GWA) for each of 23,947 measurable transcripts in the host, and 9,267 measurable transcripts in the pathogen. Unlike other eGWA studies, there was a relative absence of *cis*-eQTL that is likely explained by structural variants and allelic heterogeneity within the pathogen’s genome. This analysis identified mostly *trans*-eQTL in the pathogen with eQTL hotspots dispersed across the pathogen genome that altered the pathogen’s transcripts, the host’s transcripts, or both the pathogen and the host. Gene membership in the *trans*-eQTL hotspots suggests links to several known and many novel virulence mechanisms in the plant-pathogen interaction. Genes annotated to these hotspots provide potential targets for blocking manipulation of the host response by this ubiquitous generalist pathogen. This shows that genetic control over the co-transcriptome is polygenic, similar to the virulence outcome in the interaction of *Botrytis cinerea* on *Arabidopsis thaliana*.

## INTRODUCTION

Infectious disease is an interaction between host and pathogen. Its outcome is driven by the genetics of both organisms. The mechanisms of plant-pathogen interactions are often divided into qualitative, in which a few genetic variants of large effect shape binary disease outcomes, or quantitative, in which a spectrum of outcomes arise from the interaction of polygenic variation in the host and pathogen. The past decades have witnessed the unraveling of the molecular basis of large-effect loci on both the host side and the pathogen side that control qualitative interactions (Giraldo and Valent 2013, Marone, Russo et al. 2013, Meng and Zhang 2013, Cui, Tsuda et al. 2015, Lo Presti, Lanver et al. 2015). In this model, alternative alleles of these genes, via differential recognition events surrounding their proteins, create sweeping differences in the transcriptome and phenotypic responses to infection in both the host and pathogen. However, plant-microbe interactions cover a full range of genetic architectures, from few genes of large phenotypic effect to many genes of small effect (Poland, Balint-Kurti et al. 2009, Kou and Wang 2010, Lannou 2012). In contrast to qualitative systems, quantitative plant-pathogen interactions exhibit a lack of virulence/ resistance genes that explain large proportions of the variance in virulence in the population (Poland, Balint-Kurti et al. 2009, Kou and Wang 2010, St. Clair 2010, Roux, Voisin et al. 2014). Rather, the genetic basis of disease progression for both organisms in these interactions is highly polygenic with genetic variation influencing loci that alter a diverse array of molecular mechanisms, extending well beyond perception events (Glazebrook 2005, Nomura, Melotto et al. 2005, Goss and Bergelson 2006, Rowe and Kliebenstein 2008, Barrett, Kniskern et al. 2009, Corwin, Copeland et al. 2016, Bartoli and Roux 2017, Wu, Sakthikumar et al. 2017, Atwell, Corwin et al. 2018, Fordyce, Soltis et al. 2018, Soltis, Atwell et al. 2019). It is, however, unclear how these polygenic molecular systems in different organisms interact to alter higher-order phenotypes such as virulence, or even more direct phenotypes like the transcriptome of both species. There is some conflicting evidence on the balance of the system, with some studies and traits suggesting that genetic variation in the pathogen dominates the system (Bartha, McLaren et al. 2017, Wang, Roux et al. 2018), while others suggest a balanced contribution of plant and pathogen genetics (Corwin, Copeland et al. 2016, Fordyce, Soltis et al. 2018, Soltis, Atwell et al. 2019). Thus, there is a need to develop genomic approaches to understand how polygenic information is transmitted between the pathogen and the host to shift the genomic response of both organisms.

The polygenic variation in the pathogen should influence numerous genes that consequently shift the pathogen’s transcriptome and cause differential expression of various virulence mechanisms. This variation in virulence mechanism will then impact the host and lead to shifts in the host’s resistance-associated transcriptome. Thus, by measuring the transcriptome in both the pathogen and the host, it should be possible to map how genetic variation in the pathogen is conveyed through the pathogen’s transcriptome and concurrently how the host’s transcriptome responds. Recent work has shown that it is possible to measure the pathogen’s transcriptome in planta in *A. thaliana* - *Pseudomonas syringae* leading to new hypothesis about virulence (Nobori, Velásquez et al. 2018). In the *A. thaliana* - *B. cinerea* system, the genetic interactions are dominated by complex small-effect loci that display a high degree of interaction between the host and pathogen (Denby, Kumar et al. 2004, Rowe and Kliebenstein 2008, Zhang, Corwin et al. 2017, Atwell, Corwin et al. 2018). In this pathosystem, a co-transcriptome study with simultaneous analysis of the host’s and pathogen’s transcripts was recently done through single sample RNA-Seq (Zhang, Corwin et al. 2017, Zhang, Corwin et al. 2018). This co-transcriptome approach allowed mapping of key virulence networks in the pathogen and resistance responses within the host (Zhang, Corwin et al. 2017, Zhang, Corwin et al. 2018). Further, this study revealed a single network of transcripts from both species pathogen and host (Zhang, Corwin et al. 2018). However, these studies did not assess the genetic architecture behind these co-transcriptome interactions.

These connections can be disentangled by utilizing GWA to identify expression quantitative trait loci (eQTL)—SNPs correlated with variation in transcript expression profiles. These SNPs tag polymorphisms that cause the differential transcript accumulation and can be parsed into either *cis* or *trans* effects. Locally acting (*cis*) eQTL indicate regulatory variation within or near the expressed gene itself. *trans*-eQTL indicate SNPs that are acting at a distance and are polymorphisms that affect regulatory processes influencing the expression of the transcript. If a *trans*-eQTL affects many transcripts, it is classified as a hotspot and the SNP may influence a regulatory process that in turn influences a large number of transcripts. eQTL analysis has been utilized to study host-pathogen interactions, albeit with a focus either on host or pathogen. Frequently, these studies focus on the host’s response, such as mapping how host loci control host gene expression over time using either traditional QTL mapping or GWA analysis (Chen, Hackett et al. 2010, Hsu and Smith 2012, Zou, Chai et al. 2012, Allen, Carrasquillo et al. 2016, Christie, Myburg et al. 2017). Additional studies have begun to invert this scheme by looking at how genetic variation in the pathogen influences the host transcriptome to identify pathogen loci modulating host expression levels, and thus to identify candidate loci for interspecific signals (Wu, Cai et al. 2015, Guo, Fudali et al. 2017). These studies show that it is possible to identify pathogen loci that influence host gene expression, but they have thus far addressed pathogen populations with limited genetic variation, and thus identify the few polymorphic loci between strains with strongest effects on transcriptomic variation (Wu, Cai et al. 2015, Guo, Fudali et al. 2017). Expanding these approaches would require conducting a co-transcriptome analysis wherein both the host and pathogen transcriptomes are measured using a natural population of the pathogen, to identify a broader sampling of the pathogen loci affecting expression during infection of the host.

Thus, we conducted a GWA analysis of the pathogen and host transcriptomes to identify loci in *B. cinerea* that may be modulating the outcome of the interaction between the species. We utilized a previous co-transcriptome dataset of variation in individual transcript expression profiles of diverse *B. cinerea* isolates infecting the wildtype host Col-0 *A. thaliana* (Zhang, Corwin et al. 2017, Zhang, Corwin et al. 2018). The genomes of both the host and the pathogen harbor extensive genetic diversity that has been successfully used for GWA to identify loci controlling virulence (Atwell, Corwin et al. 2018, Soltis, Atwell et al. 2019). Further, the virulence outcome of the interaction is easily measured via high-throughput digital imaging and has previously been utilized for studies into the transcriptomic and genomic basis for virulence allowing for a large body of molecular information to underpin any hypothesis generation from GWA (Denby, Kumar et al. 2004, Rowe and Kliebenstein 2008, Zhang, Corwin et al. 2017, Atwell, Corwin et al. 2018, Soltis, Atwell et al. 2019). The loci tagged by these SNPs have an explicit directionality of effect, as genetic causality must arise within the pathogen and then extend to the host. Our analysis found mostly small-effect polymorphisms dispersed throughout the *B. cinerea* genome, with several *trans*-eQTL hotspots. These hotspot loci are linked to specific host or pathogen transcript co-expression modules and to variation in lesion size. There was no identifiable overlap in the hotspots that influenced the host’s or the pathogen’s transcriptome, suggesting a surprisingly independent basis of transcriptional regulation of host and pathogen by the *B. cinerea* genome. Among these hotspot loci, all appeared to tag novel genes not previously identified as controlling plant-pathogen virulence interactions. Overall, we identify a mix of novel loci potentially controlling the interaction of *A. thaliana* and *B. cinerea* via modulation of gene expression, with evidence for connections to virulence.

## RESULTS

### eQTL indicate polygenic transcriptome modulation

To better understand how natural genetic variation in the pathogen influences both the host and pathogen transcriptomes, we performed expression GWA across all genes expressed in both species within the *B. cinerea* - *A. thaliana* pathosystem. This incorporated the expression profiles of 9,267 *B. cinerea* genes and 23,947 Col-0 *A. thaliana* genes, each as individual traits across 96 diverse *B. cinerea* isolates. For each trait, we used a Genome-wide Efficient Mixed Model Association (GEMMA) model with a previous *B. cinerea* genome-wide SNP dataset of 237,878 SNPs with a conservative minimum minor allele frequency of 0.20 (Zhou and Stephens 2012, Atwell, Corwin et al. 2018). GEMMA estimates the significance of effects of each SNP on the focal trait as a p-value after accounting for potential effects of population structure within the *B. cinerea* isolates. In total, GEMMA was able to identify *B. cinerea* SNPs linked to transcriptional variation in 5,213 *A. thaliana* genes and 1,616 *B. cinerea* genes (Figure 1). For these genes with significant SNPs, there was a median of 10±25 SNPs per transcript for *B. cinerea*, and a median of 10±13 SNPs per transcript or *A. thaliana* transcripts (Figure S1a, S1b). Filtering the SNP-transcript associations for smallest p-value or largest effect estimate, we detected no *cis*-effect polymorphisms in the top 50 candidates (Supplemental Data Set 4). Further, the distribution of p-values for significant SNPs suggest a highly polygenic basis of loci modulating transcriptome variation (Figure S1c, S1d, Figure S2).

**Figure 1.**
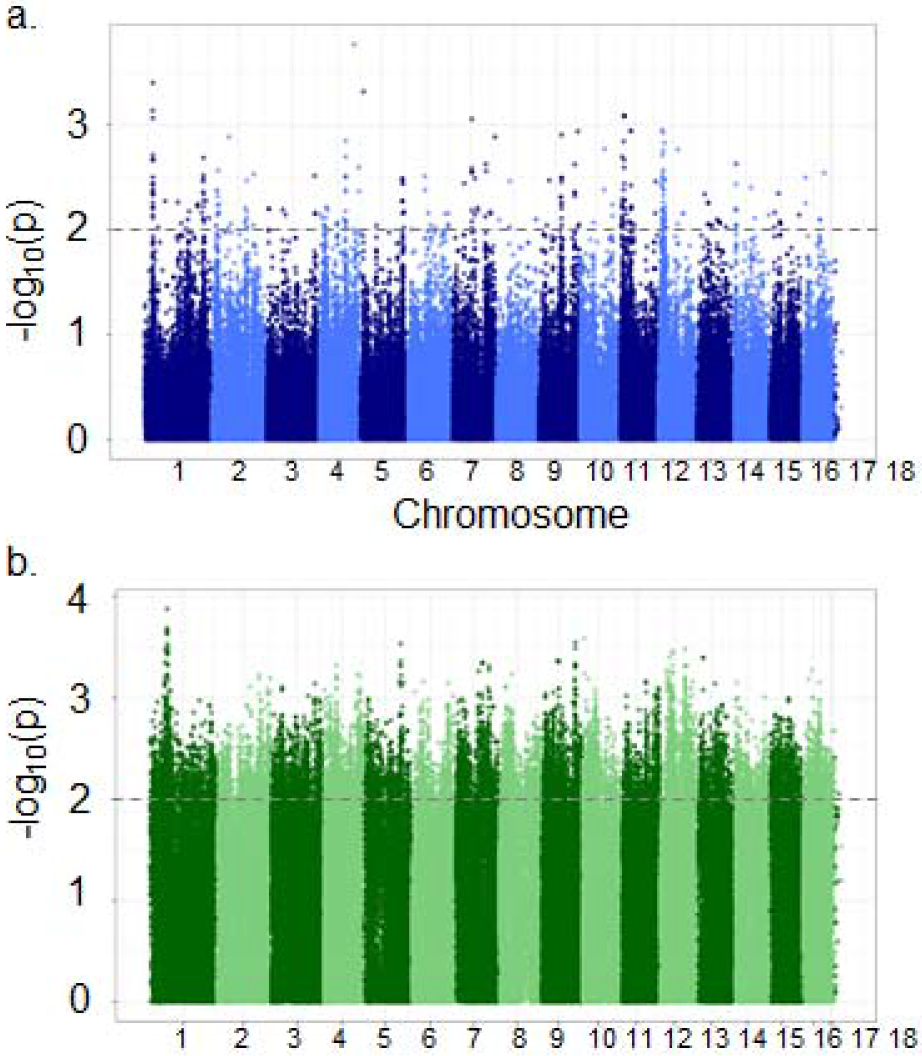
Manhattan plot examples for one transcript per species. Panel a is an example plot of p-values for all *B. cinerea* SNP associations to a single *B. cinerea* transcript, from Bcin01g00170. Panel b is an example plot of p-values for all *B. cinerea* SNP associations to a single *A. thaliana* transcript, from AT1G01010. hed line indicates significance at p = 0.01.

Given the scale of this dataset at 33,214 transcript phenotypes, it was not viable to estimate empirical significance thresholds for every transcript using 1,000 or more permutations. However, we permuted the whole dataset across each of the tens of thousands of traits five times to break all real phenotype-genotype associations and repeated GEMMA to get a feel for the potential for dominant patterns that may exist randomly (individual expression profiles in *B. cinerea* and *A. thaliana*). We then compared the permuted minimum p-value per transcript across all SNPs to the data obtained from real traits. This showed that the top SNP per trait for most genes show a stronger association in our observed data than across any of the 5 permutations. In *B. cinerea*, the observed p-value is lower for 69% of genes, much more than the expected 17% due to random chance, and in *A. thaliana* the observed p-value is lower for 58% of genes. Thus, to develop genomic summary images of the results, we focused on the top significant SNP per transcript for the remaining analysis. This allows us to focus on SNPs that are highly likely to be associated with the trait. To account for the potential that the top SNP may not fully encompass the genomic signal, we also assessed any general pattern using the top 10 SNPs.

### Absence of transcriptome cis-effect dominance

A hallmark of eQTL mapping studies using either GWA or structured mapping populations in a wide range of species is the occurrence of large-effect loci that map to the gene itself, i.e. *cis*-eQTL or *cis*-SNPs (Brem, Yvert et al. 2002, Schadt, Monks et al. 2003, Monks, Leonardson et al. 2004, Keurentjes, Fu et al. 2007, West, Kim et al. 2007, Zou, Chai et al. 2012). To test if the *B. cinerea* transcriptome shows a similar *cis*-eQTL pattern, we plotted the position of the transcript’s genomic position against the top GWA SNP for all the *B. cinerea* transcripts. We first focused on the single top SNP hit per transcript, with the lowest p-value (strongest evidence of significant effect on expression) in the gene of interest. If control of gene expression is localized to the gene itself or to proximate loci, we would expect a strong linear (*cis*-diagonal) association between the center of each gene and the genomic location of its top SNP hit. However, there was no evidence of any cis-diagonal (Figure 2). This pattern held whether we examine the top SNP per transcript (Figure 2a) or the top 10 SNPs per transcript (Figure 2b). In contrast, there was evidence for *trans*-eQTL hotspots; loci which modulate expression variation across many of the pathogen genes (Figure 2).

**Figure 2.**
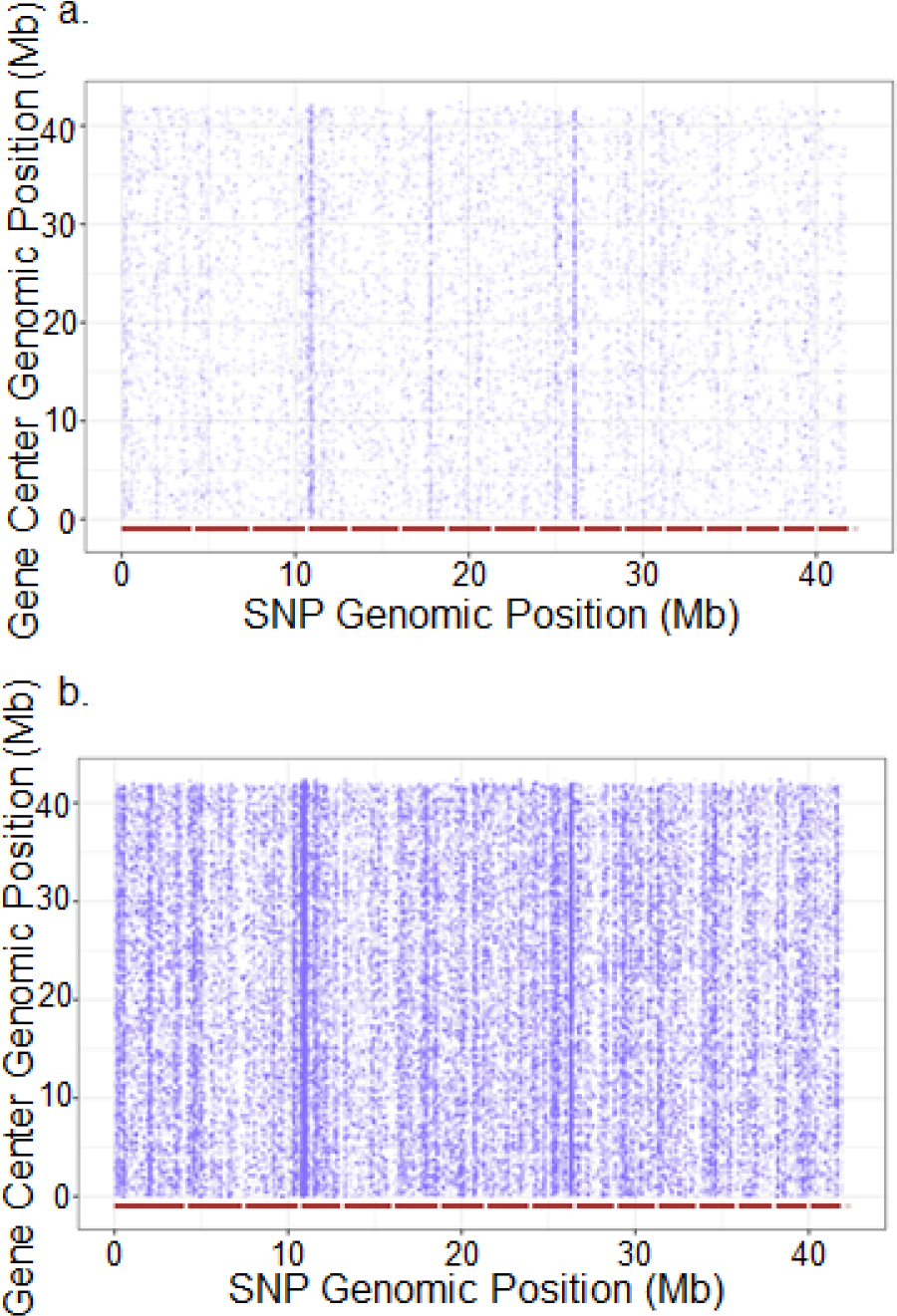
*cis*-diagonal plot comparing *B. cinerea* gene center to position of top associated SNP, for all 9,284 transcripts mapped to the *B. cinerea* genome. We retained only the SNPs with highest probability (lowest p-value) of significant effect on expression for each transcript. Panel a depicts the single top SNP per transcript. Panel b depicts the top 10 SNPs per transcript. The *18 B. cinerea* chromosomes are delimited by red bars along the x-axis, and positions indicate individual SNPs. The y-axis depicts the same chromosome alignment, but positions are the center of each mapped transcript. Vertical striping of SNP positions indicates genomic locations of putative *trans*-eQTL hotspots.

To test if there might be a bias towards *cis*-effects that may function at a close distance, we calculated the distance between the center of each transcript and the top associated SNP. If *cis*-acting loci contribute the bulk of genetic control of expression variation, we would expect to see a high frequency of short-distance associations, and a rapid decline to a plateau moving away from the gene of interest. However, we observe that distances between transcript center and top SNP as far as 2 Mb are common (Figure S3). These distances are similar to what would happen if the causal SNPs had no *cis*-association and were instead scattered across the genome (Figure S3). We further investigated with 50 SNPs with the smallest p-value of association and largest effect size estimates as these are frequently assumed to be in *cis*. However, none of these SNPs were in *cis* to the transcript being tested, further supporting the absence of a dominant pattern of *cis*-effects (Supplemental Data Set 4). As such, we do not see evidence for *cis*-effect loci overrepresented in the top candidates for control of expression variation. Rather, most of the loci that we can associate with potentially influencing gene expression variation in *B. cinerea* on *A. thaliana* is *trans*-acting.

### Search for *cis*-effects through focus on gene networks with presence-absence polymorphism

The absence of a dominant *cis*-pattern in the genome-wide transcript-to-SNP associations could be caused by a relative absence of *cis*-variation. Alternatively, haplotype heterogeneity or allele frequency may complicate the ability to accurately identify *cis*-polymorphisms (Chan, Rowe et al. 2010, Rivas, Beaudoin et al. 2011, Visscher, Wray et al. 2017). To test between these possibilities, we conducted a more focused analysis on three biosynthetic pathways that exist as gene clusters: the botcinic acid biosynthetic pathway (13 genes, 55.8 kb), botrydial biosynthetic pathway (7 genes, 26 kb), and a putative cyclic peptide pathway (10 genes, 46.5 kb) (Deighton, Muckenschnabel et al. 2001, Colmenares, Aleu et al. 2002, Porquier, Morgant et al. 2016, Zhang, Corwin et al. 2018). These pathways have known presence-absence polymorphisms segregating in this population and should have *cis*-eQTL but none were detected by our analysis (Siewers, Viaud et al. 2005, Pinedo, Wang et al. 2008, Zhang, Corwin et al. 2018). Genes of the botcinic acid biosynthetic pathway and the botrydial pathway are variable across the *Botrytis* genus as well (Valero-Jiménez, Veloso et al. 2019). Critically, the transcripts within each of these pathways are highly correlated across the isolates, suggesting that their expression variation is controlled by pathway-specific variation (Zhang, Corwin et al. 2018). Thus, these loci may have false-negative issues that prevented the detection of real *cis*-eQTL.

To test if these pathways have undetected *cis*-eQTL we used all of the SNPs for each biosynthetic cluster to investigate haplotype diversity across the *B. cinerea* isolates. We first investigated the botcinic acid cluster which identified a number of distinct haplotypes with a few individual outlier isolates (e.g. B05.10, Fd1) (Figure 3a). We then utilized the haplotypes to test for specific effects on transcript expression for the biosynthetic pathway. This identified a single clade of isolates with a distinctly lower level of expression than the other clusters (Figure 3b). Investigating the short-reads and SNP calls showed that these 12 isolates share a 53.5 kb deletion that removes the entire biosynthetic cluster, from Bcboa1 to Bcboa17 (Figure 3c). After removing the major deletion, we found no remaining significant effect of cluster membership in the remaining 3 major clusters on expression profile (F(1,74)=0.36, p=0.55). However, within each of these clusters there are independent isolates with very low pathway expression, suggesting loss-of-expression polymorphisms (Noble Rot, 01.04.03, Apple 517, 02.04.09) (Figure 3b). While these isolates each contain smaller deletions that are independent of each other, it is not clear what is functionally leading to the loss of botcinic acid biosynthetic pathway expression (Figure 3c). This suggests that for this clustered pathway, there are undetected *cis*-effect polymorphisms, a large common deletion and rarer additional events. In-depth analysis of the botcinic acid biosynthetic cluster has thus far identified one transcription factor (Bcboa13) that controls expression of the cluster (Porquier, Moraga et al. 2019). However, none of the SNPs within or near Bcboa13 were significantly associated with variation in expression of the botcinic acid biosynthesis genes.

**Figure 3.**
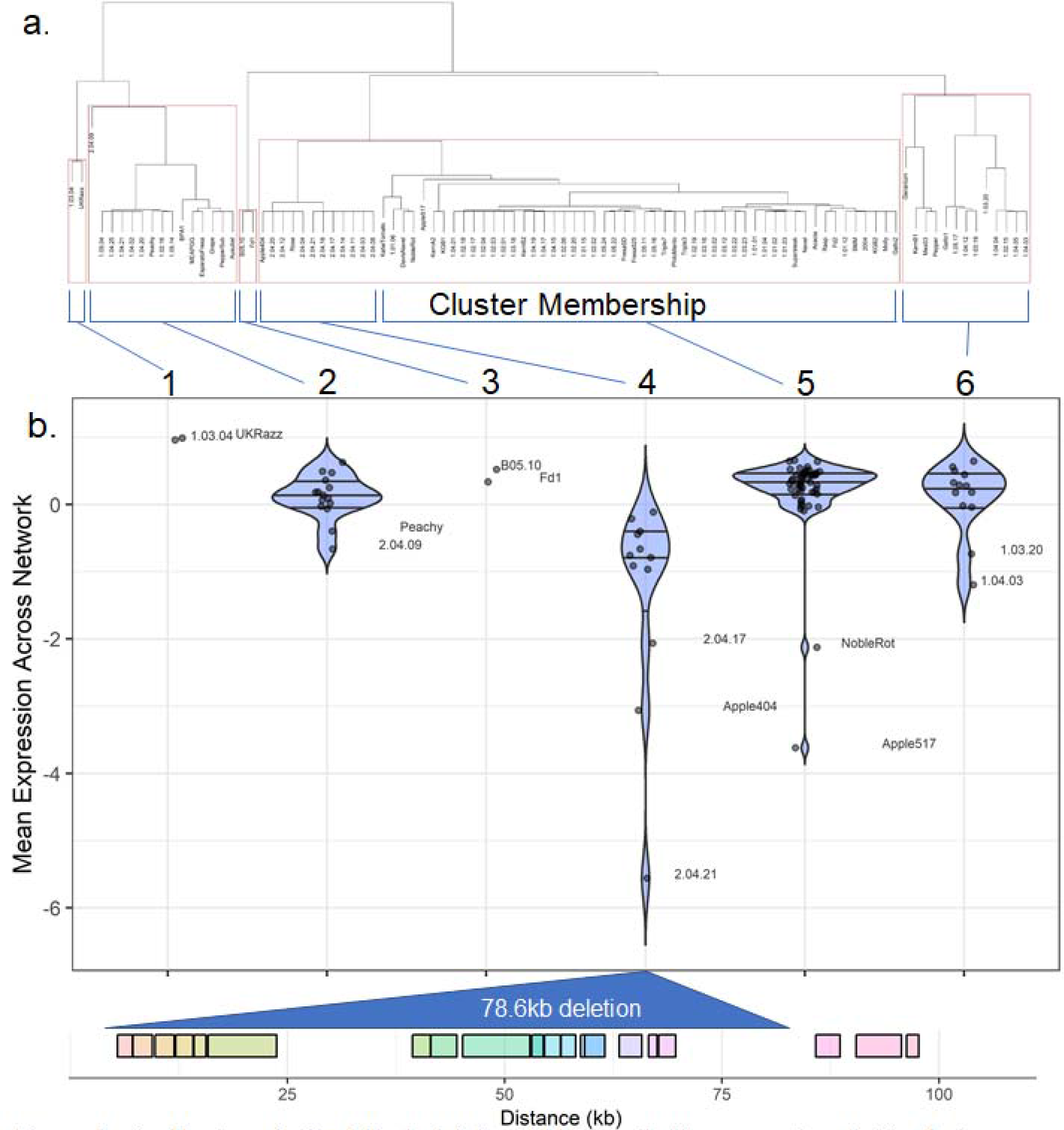
*cis*-effect analysis of the botcinic acid biosynthetic gene network. Panel a is hierarchical clustering of *B. cinerea* isolates from SNPs within the botcinic acid biosynthetic gene network. Clustering was based on mean linkage (UPGMA), with correlation distance and 1000 bootstrap replications. AU p-values are reported in red, BP values in green. Edges with high AU values are considered strongly supported by the data, and clustering is drawn according to these edges with AU > 95%. Panel b is Violin plots of botcinic acid network-level expression within *B. cinerea* clusters. Isolates are clustered based membership in groups defined by hierarchical clustering of the SNPs within the botcinic acid biosynthesis network. Panel c is the gene models of the biosynthetic gene network, with each box delimiting a single gene (including Bcboa1 to Bcboa13 and 5 additional genes) and with the cluster 3 deletion indicated as a triangle.

**Figure 4.**
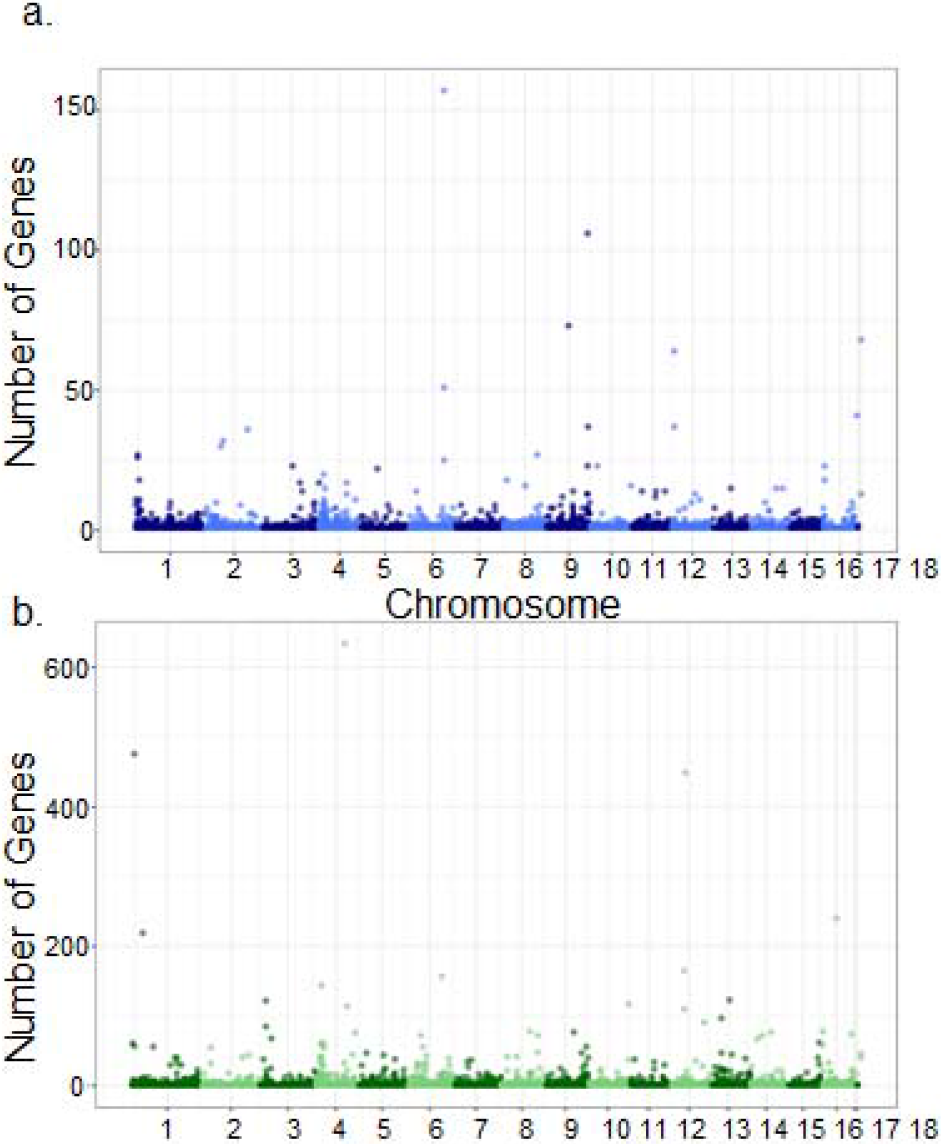
Manhattan-type plot of GEMMA results of transcriptome-wide *B. cinerea* and *A. thaliana* expression phenotypes. Panel a is a Manhattan-type plot of the top 1 SNP hit per *B. cinerea* transcript on Col-0 *A. thaliana*. Panel b is a Manhattan-type plot of the top 1 SNP hit per *A. thaliana* transcript when infected by *B. cinerea*.

We then investigated the other two biosynthetic pathways for additional evidence of missed *cis*-acting genetic variation. The botrydial biosynthetic network, and the cyclic peptide pathway, exhibit a lack of dominant *cis*-effect SNP patterns much like the botcinic acid biosynthetic network. Hierarchical clustering within each of these networks by genic SNP variation divides the isolate population into two groups that are not associated with mean pathway expression (Figure S4, Figure S5). However, within the cyclic peptide pathway, minor deletions within the intergenic regions correlate with low expression, and two isolates with partial deletions within the genes early in the pathway exhibit very low pathway expression (1.05.16, 1.05.22) (Figure S5). In contrast, there was no evidence for SNP *cis*-effects, and this pathway did not harbor any obvious loss-of-expression events (Figure S5). As such, we can detect *cis*-acting variation in the form of deletions for two of the biosynthetic pathways. This suggests that there are missing *cis*-effects within the *B. cinerea* GWA, likely missed due to the frequency of structural variants often falling below the minor allele cutoffs. Testing whether insertion and deletion events account for the majority of localized control of expression variation would require long-read sequencing to accurately identify these structural variants and computational approaches that can blend SNP and indel information.

### Detection of *trans*-eQTL hotspots

While *cis*-effects are difficult to identify, there was a strong signature of SNPs that appeared to affect more transcripts than expected by chance (Figure 2). These are considered positions where there is a causal polymorphism that influences the regulation of numerous genes in *trans*, i.e. a *trans*-eQTL hotspot. In this dataset, we can extend this analysis to look for *trans*-eQTL hotspots that extend beyond *B. cinerea* and influence the expression of genes in the host, *A. thaliana*. We queried for hotspots in both the *B. cinerea* and *A. thaliana* transcriptome by using overlaps in the top SNP per transcript (Figure 4, Figure 5). By permuting the SNP positions, we identified maximum permuted hotspot sizes as a SNP associated with 11 *B. cinerea* transcripts or 80 *A. thaliana* transcripts. For further analysis of hotspots, we utilized a conservative threshold of 20 linked transcripts for *B. cinerea* and 150 transcripts for *A. thaliana*. This analysis identified 13 SNPs as potential *trans*-eQTL hotspots for the *B. cinerea* transcriptome and 12 SNPs as potential cross-species *trans*-eQTL influencing *the A. thaliana* transcriptome (Figure 5, Figure 6). The *trans*-eQTL hotspots are spread throughout the genome (Figure 6, Table 1).

**Table 1.** Annotation of the hotspots identified from *B. cinerea* and *A. thaliana* eQTL. Each row identifies a significant eQTL hotspot SNP associated with transcripts in *B. cinerea* or in *A. thaliana*. Gene functions are from BotPortal, Arabidopsis GO overrepresentation are from PANTHER. The table also presents if the gene’s transcript is correlated with virulence or is associated with virulence via GWA from previous analysis (Zhang, Corwin et al. 2017, Atwell, Corwin et al. 2018, Zhang, Corwin et al. 2018).

| hotspot gene | hotspot SNP | Botrytis Transcripts | Arabidopsis Transcripts | Name | gene function | Virulence Correlation | Virulence GWA | Arabidopsis GO overrepresentation |
| --- | --- | --- | --- | --- | --- | --- | --- | --- |
| Bcin01g01610 | 629157 | 0 | 219 |  | Glucose/ribitol dehydrogenase | NA |  | amylopectin, glycogen, chlorophyll, chloroplast, photosynthesis |
| Bcin02g02480 | 910858 | 31 | 6 |  |  | p=0.008 |  | NA |
| Bcin02g02850 | 1032242 | 32 | 0 |  | Fructosamine-3-kinase |  |  | NA |
| Bcin03g00960 | 336467 | 1 | 122 |  | GTP cyclohydrolase I | NA |  | carotenoid biosynthesis, chloroplast organization |
| Bcin03g05020 | 1695103 | 24 | 15 |  |  | NA |  | NA |
| Bcin04g00830 | 310509 | 0 | 144 |  | NACHT nucleoside triphosphatase |  |  | nucleic acid metabolism |
| Bcin04g04700 | 1633072 | 0 | 634 |  | Heterokaryon incompatibility | NA |  | photosynthesis, light, translation |
| Bcin04g05160 | 1791561 | 3 | 114 |  |  | NA |  | chitin response |
| Bcin05g02780 | 1015184 | 22 | 2 |  |  | NA |  | NA |
| Bcin06g05680 | 1952544 | 0 | 157 |  | WLM | p < 0.001 |  |  |
| Bcin06g05790 | 1988179 | 25 | 0 |  |  | NA |  | NA |
| Bcin08g05340 | 2014228 | 27 | 0 |  | F-box domain | NA |  | NA |
| Bcin09g03390 | 1235841 | 73 | 6 |  | 6-phosphogluconate dehydrogenase | NA |  | NA |
| Bcin09g06590 | 2330312 | 106 | 56 |  | SNF2-related; Helicase, superfamily 1/2 | NA |  | metabolism |
| Bcin09g06590 | 2334368 | 23 | 0 |  | SNF2-related; Helicase, superfamily 1/2 |  |  | NA |
| Bcin10g00940 | 383007 | 23 | 1 |  |  | p < 0.001 |  | NA |
| Bcin10g05900 | 2268522 | 0 | 117 |  | Winged helix-turn-helix Transcription factor |  |  | water stress |
| Bcin12g00330 | 115491 | 37 | 1 | Bcpat1 | Topoisomerase II-associated protein PAT1 | p < 0.001 |  | NA |
| Bcin12g00330 | 115511 | 64 | 3 | Bcpat1 | Topoisomerase II-associated protein PAT1 | p < 0.001 |  | NA |
| Bcin12g02130 | 758420 | 0 | 110 |  |  | NA |  | response to stimulus |
| Bcin12g02130 | 760499 | 2 | 265 |  |  |  |  | JA, fungal response, microbe defense, biotic stress |
| Bcin12g02340 | 842369 | 1 | 449 | Bccds1 | Phosphatidate cytidyltransferase | p < 0.001 |  | primary metabolism, amino acid biosynthesis, salt stress, biotic response |
| Bcin13g02930 | 1026752 | 0 | 123 |  | SET domain | NA |  | transport |
| Bcin16g00010 | 76596 | 23 | 2 |  | SsuA/THI5-like | NA | Yes | NA |
| Bcin16g01950 | 787512 | 0 | 240 | Bccwh41 | Glycoside hydrolase, family 63 | NA |  | photosynthesis |

**Figure 5.**
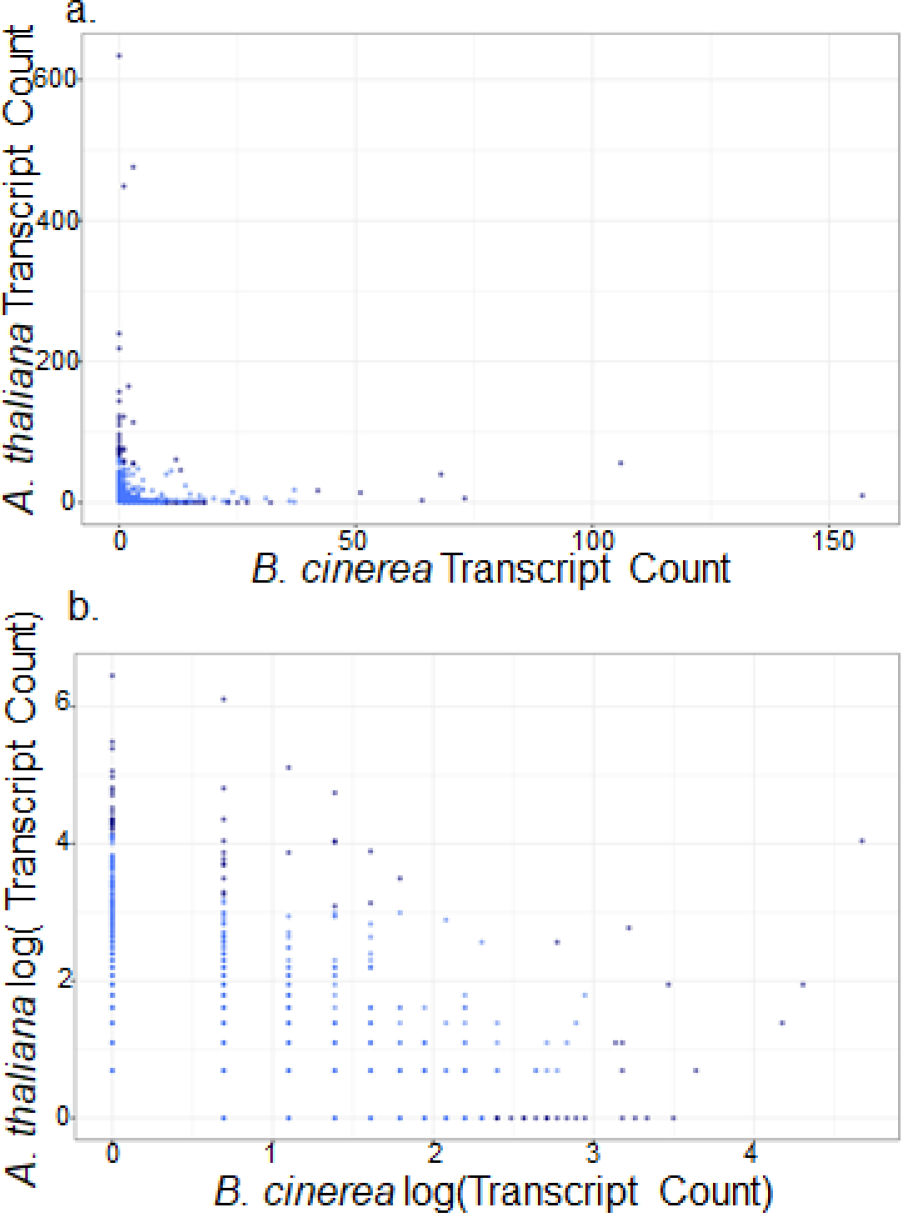
Interspecific hotspot comparison on the *B. cinerea* genome. For each SNP that is a top hit for one or more transcripts, the number of associated transcripts is counted, across both the *B. cinerea* transcriptome and the *A. thaliana* transcriptome.

**Figure 6.**
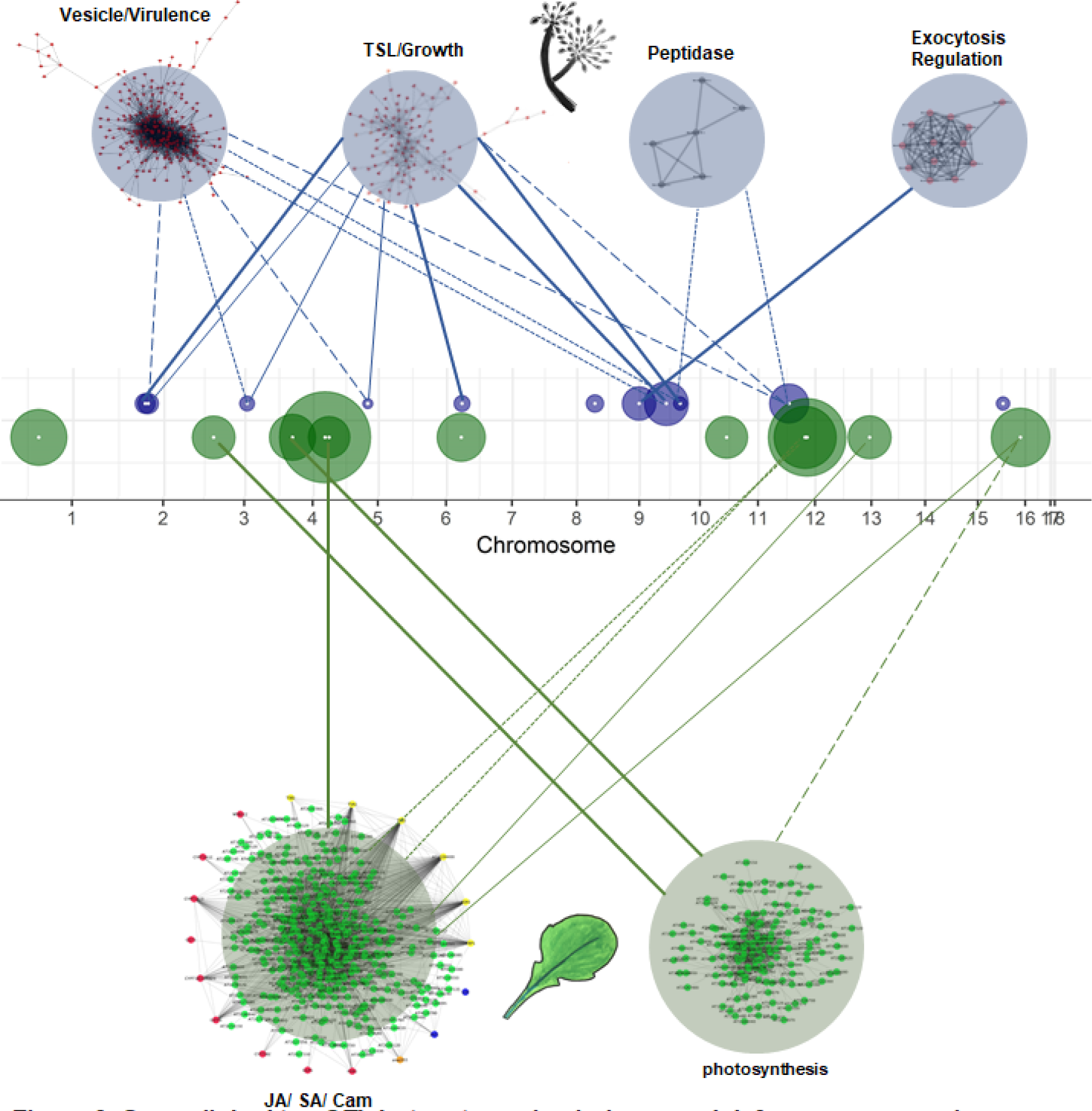
Genes linked to eQTL hotspots are in virulence and defense co-expression networks. Circles along the *B. cinerea* 18 chromosomes are eQTL hotspots, centered at the gene containing the eQTL and with radius proportional to the number of transcripts linked to this hotspot. The gene center is marked with a white dot. Hotspots for *B. cinerea* transcripts are drawn in blue, hotspots for *A. thaliana* transcripts are drawn in green. The network names are based on biological functions from gene ontology analysis of network members, from Figure 4 of Zhang *et al*. 2018 and Figure 6 of Zhang *et al*. 2017. The *A. thaliana* networks depicted are the most inclusive of the host-dependent networks, from *npr1-1*. Links between hotspots and co-expression networks are drawn according to the number of genes shared between them. Variable line weight represents the percent of hotspot target genes shared with the co-expression network; 1-25% is dashed, 25-50% is dotted, 50-75% is solid, 75-100% is heavy solid.

The benefit of a co-transcriptome is that it should be possible to map how polymorphisms cause effects in the pathogen and how these effects transmit to an altered transcriptome in the host. This would suggest that a *trans*-eQTL hotspot for *B. cinerea* transcripts may control virulence pathways and thus cause an associated *trans*-eQTL hotspot in the *A. thaliana* response. However, through the GWA analysis we detected no significant overlap in eQTL hotspots across the two transcriptomes; hotspots targeting *B. cinerea* gene expression linked to only 0 to 56 transcripts in *A. thaliana*, and hotspots targeting *A. thaliana* gene expression linked to only 0 to 3 *B. cinerea* transcripts. All of these are values that are below the permutation threshold. To identify overlap in these eQTL hotspots missed by GWA due to low power in the single top SNP per transcript, we repeated the full analysis by selecting the top 10 SNPs per transcript. This again identified a limited number of *trans*-eQTL hotspots with little overlap between the two species’ transcriptomes (Figure S6). This suggests that the pathogen’s influence on the host’s transcriptome is not solely limited to major interactions between *tran*s-eQTL hotspots but can involve narrower changes in the pathogen that are magnified in the host’s response. One possible explanation for this is if a SNP alters the expression of a single effector gene or mechanism in the pathogen that has no impact on the pathogen but impacts the host. For example, altering expression of the botcinic acid biosynthetic cluster would alter the accumulation of the metabolite that then caused large responses in the host. In addition, it is possible that some of this low overlap could be caused by the high level of variation in this pathogen causing a drop in power to detect these overlaps. Deeper analysis into the transcriptome and downstream responses could elucidate how restricted responses in the pathogen transcriptome translate to sweeping responses in the host. However, future studies using these eQTL hotspots as *a priori* candidates for control of transcript variation in both host and pathogen may increase power to detect more modulation overlap across the two transcriptomes.

### eQTL hotspot modules

To better understand the gene expression modules that are being influenced by these hotspots, we examined the genes influenced by each hotspot. We first utilized the gene ontology (GO) annotations within each species to better assess if there was any common functionality. The *B. cinerea* GO annotations showed a preponderance of enzyme, signal peptides for secretion, and transcription factor annotations but no specific molecular insights arose largely because the majority of genes had no annotation (Table 1, Supplemental Table 1, Supplemental Data Set 1, Supplemental Data Set 4). In contrast, GO analysis of the *A. thaliana* transcripts showed that three of the hotspots have an overrepresentation of photosynthesis-related functions within their targeted genes (Table 1, Supplemental Table 3, Supplemental Table 4). Downregulation of photosynthesis gene expression is a hallmark of plant immune processes (Bilgin, Zavala et al. 2010, Jiang, He et al. 2017). Two of the hotspots predominantly affect *A. thaliana* genes associated with abiotic stress responses. Only two of the hotspots are directly linked to expected plant defense loci, including chitin response and microbe defenses. This suggests that the *B. cinerea* genes underlying these hotspots have specific effects on defined networks within the host and are not causing nonspecific responses.

In previous work, we had defined key transcript modules within both the host and pathogen transcriptomes that could be linked to virulence (Jiang, He et al. 2017, Zhang, Corwin et al. 2018). Thus we proceeded to test if any of these *trans*-eQTL hotspots were associated with the previously defined transcript modules, by comparing gene lists for overlap between module membership and hotspot association (Zhang, Corwin et al. 2017, Atwell, Corwin et al. 2018, Zhang, Corwin et al. 2018) (Table 1). Nine of the 11 *B. cinerea* eQTL hotspots were enriched for transcripts present in one or more of four major *B. cinerea* co-expression networks on *A. thaliana* (Figure 6). An additional six *B. cinerea* co-expression networks did not share any gene membership with our eQTL hotspots. In particular, two of the eQTL-enriched networks were host-specific networks functionally associated with virulence, with 7 of the 11 *B. cinerea* hotspot genes associated with one of these virulence co-expression networks. Similarly, nine of the *A. thaliana* eQTL hotspots were enriched for transcripts from two of the major *A. thaliana* co-expression networks when infected with *B. cinerea* (Figure 6). These two modules contain genes that function in jasmonate and salicylic acid signaling processes and camalexin biosynthesis (Network I), or photosynthesis (Network IV). Interestingly, these links are not limited to a single hotspot, but are strong connections across a number of different hotspots suggesting that these modules have a polygenic architecture underlying them (Figure 6). These frequent links suggest that the identified eQTL hotspots may exhibit regulatory control over co-expressed modules of genes active in virulence interactions between *B. cinerea* and its host. If these eQTL hotspots are modulating expression of many genes, and affecting lesion size, they may be major *B. cinerea* control points in the plant-pathogen interaction.

### eQTL hotspot candidate genes

To better understand what the causal basis of these hotspots might be, we investigated the candidate genes associated with the SNPs. The 12 *B. cinerea* hotspots linked to *A. thaliana* transcripts, annotated to 11 genes, included 4 enzymes and 2 genes associated with isolate compatibility (Table 1). The 13 *B. cinerea* hotspots linked to *B. cinerea* expression profiles were associated to 11 genes, including 4 enzymes (Table 1). However, only one of these 22 genes had any previous published information linking them to virulence functions in *B. cinerea* or other fungi; a glycoside hydrolase whose homolog shows increased expression in virulent strains of *Ustilago maydis* on *A. thaliana* (Bccwh41) (Martínez-Soto, Robledo-Briones et al. 2013). To test if any of these 22 eQTL hotspot genes may have a link with virulence in *B. cinerea*, we compared their expression in the co-transcriptome data to existing virulence measurements. The virulence was measured on different leaves, and not on the same leaves as the co-transcriptome. Transcript accumulation for three *B. cinerea* hotspot genes and two of the *A. thaliana* hotspot genes are strongly positively correlated to lesion size variation, and none are negatively correlated with lesion size (Table 1) (Zhang, Corwin et al. 2018). Further, we utilized a previous GWA of virulence of these same isolates on *A. thaliana* to test if there was any overlap. This showed that one of the *B. cinerea* hotspot genes (Bcin16g00010, SsuA/THI5-like) is a top GWA hit controlling lesion size across host genotypes and association methods (Table 1) (Atwell, Corwin et al. 2018). Together, this suggests that these genes are likely candidates for controlling transcriptome responses in both the host and pathogen.

## DISCUSSION

### Complications in detection of *cis*-acting loci

The vast majority of eQTL studies identify a dominant signature of *cis*-acting loci. However, in *B. cinerea*, the dominant pattern was one of *trans*-eQTL with few detected *cis*-eQTL. A deeper investigation suggested that this may be due to genetic factors that complicate our ability to identify the *cis*-acting SNPs. *B. cinerea* has high haplotype diversity, and in the three gene clusters investigated, there were potential rare *cis*-acting variants that fall below the minor allele cutoff for GWA (Atwell, Corwin et al. 2018). The identified *cis*-acting variants were often deletions, which further complicate the ability to identify a *cis*-eQTL signature by introducing non-SNP variation that is missed by the GWA algorithm. This may account for additional undetected *trans*-eQTL as well. Additional cis-acting variants may best be captured by variation in transposons (Porquier, Moraga et al. 2019). To fully understand the pattern of potential *cis*-acting loci in *B. cinerea* would require a deeper investigation of structural variation by incorporating long-read sequencing. Future GWA with a larger sample of diverse pathogen isolates and deeper sequencing would assist with identifying these *cis*-eQTL. Additionally, the GWA algorithms would need to be written to allow for simultaneous use of both SNP and presence/ absence polymorphism data; one option is to code deletions as an additional state for each genotyped variant (Wang, Roux et al. 2018). This does suggest that there is likely a significant fraction of undetected *cis*-eQTLs within *B. cinerea*, caused by the high polymorphism rate within this species.

### Dispersed interactions across host and pathogen genomes

Using co-transcriptome GWA, we identified 25 *trans*-eQTL hotspots dispersed across the *B. cinerea* genome that modulate either the host or pathogen transcriptomes. This contrasts with previous cross-species eQTL studies, which identified one or only a few cross-species eQTL hotspots (Wu, Cai et al. 2015, Guo, Fudali et al. 2017). Further, most of the genetic variation detected in our study is distant from the affected transcripts, *i.e*. located in *trans*. These *trans*-eQTL hotspots are linked to the expression variation for five major *B. cinerea trans*- co-expression networks with genes dispersed across the genome (Zhang, Corwin et al. 2018). In particular, the eQTL hotspots influenced the expression of many genes from the previously identified *B. cinerea trans*-co-expression networks (vesicle/virulence, translation/growth, exocytosis regulation, peptidase). Interestingly, the candidate polymorphisms are spread throughout the genome and the detected eQTL hotspots are not in regions of the genome with outlier levels of genetic variation. Further, the genetic targets of these eQTL are dispersed across the plant and pathogen genomes (Zhang, Corwin et al. 2018). As such, *B. cinerea* does not fit the models of what might be expected in filamentous fungi that have multiple-speed genomes due to varying selective pressures across the genome. In these specialist fungi with closer co-evolution with their host species, diverse fungal virulence effectors are enriched in regions of the genome containing repetitive sequences and transposable elements (Dong, Raffaele et al. 2015). These regions show enhanced rates of mutation and polymorphism while the rest of the genome shows slower evolutionary rates. If this pattern defined variation in the current system, it would predict clustering of the great majority of eGWA hits to a few locations, rather than distribution of eQTL across the genome as was found. This is consistent with previous findings of high diversity in *B. cinerea* genome-wide, and virulence mapping to large swaths of the pathogen genome including 16 of 18 chromosomes (Atwell, Corwin et al. 2018, Caseys, Shi et al. 2018). It will require conducting a similar analysis in the multi-speed genome filamentous fungi to test whether eQTL in a pathogen with a multi-speed genome truly cluster within the highly polymorphic regions. These findings together provide evidence for polygenic *trans*-regulation of gene expression in *B. cinerea* interactions that then coalesces around specific transcriptional modules to influence virulence.

### Polygenic modules and pleiotropy in cross-species eQTL

Previous pathogen-linked eQTL studies typically identified more explicit patterns whereby each host expression profile was explained by only a single major-effect pathogen locus (Guo, Fudali et al. 2017) or each pathogen eQTL linked to a specific host network (Wu, Cai et al. 2015). In contrast, co-transcriptome eGWA with *B. cinerea* identified a more complex picture with numerous *trans*-eQTL hotspots and these linked to multiple transcriptome modules in either the host or the pathogen. This suggests that the polygenic architecture of the pathogen may at least in part function by influencing these defined modules rather than functioning as thousands of individual genes each separately targeting the host. This gives us an overarching pattern of polygenic and pleiotropic genetic regulation, as both the host and pathogen appear to draw from extensive genetic variation to determine disease outcomes. In effect, we see polygenicity of host expression regulation by the pathogen at the gene level, and at the network level. It remains to be ascertained if this is a system to create robustness in these connections in the face of changes to the pathogen or host genetics or if, alternatively, this is an indication that there are discrete set of interaction mechanisms between the host and the pathogen.

### Diverse mechanisms linked to candidate causal loci

Investigating the putative function of the candidate loci underlying the different *trans*-eQTL hotspots identified an array of potential molecular mechanisms. While one might assume that transcription factors are the most likely genes in which genetic variation would lead to *trans*-eQTL hotspots, there was instead an enrichment for enzyme-encoding genes as the identity of these loci. This included four enzymes linked to the 13 *trans*-eQTL hotspots influencing the *B. cinerea* transcriptome and an additional four linked to the 12 *trans*-eQTL hotspots influencing the *A. thaliana* transcriptome. Interestingly, these enzymes were largely linked to various aspects of sugar release from the plant cell wall, or potential reactions involving sugar-phosphates (Table 1). In addition to enzymes, the candidate genes for four of the *trans*-eQTL linked to transcriptional regulators. While one, Bcin10g05900, a putative winged helix TF, would be predicted to potentially have pathway specific effects, the other three were more likely to have general effects on transcription including Bcin12g00330, a putative Topoisomerase II-associated protein PAT1, and Bcin09g06590, a putative helicase (Table 1). Interestingly, the putative winged helix TF was linked to a trans-eQTL hotspot influencing the expression of *A. thaliana* genes that show an over-representation of genes linked to water deprivation. This suggests that this putative winged helix TF may influence a specific virulence factor that influences this Arabidopsis network. Interestingly, while the candidate genes are linked to processes that likely influence virulence, none of them have been explicitly shown to influence virulence in *B. cinerea*. Future work is necessary to begin testing these loci and if and how they may influence virulence and the host/pathogen co-transcriptome.

## Conclusion

Previous work in the *B. cinerea* – *A. thaliana* pathosystem established connections between host polymorphisms and lesion growth, between gene expression and lesion size, and between transcriptomes of the host and pathogen (Corwin, Subedy et al. 2016, Zhang, Corwin et al. 2017, Fordyce, Soltis et al. 2018, Zhang, Corwin et al. 2018). This study begins to establish the foundation to begin testing directional causal inferences from pathogen genome to transcriptome to disease phenotype by connecting genetic variation in the pathogen to expression changes in both the host’s and pathogen’s transcriptomes. This showed a preponderance of *trans*-acting polymorphisms with predominantly moderate to small effects, suggesting that a polygenic architecture underlies the transcriptome variation, similar to the virulence interaction. Using previously defined transcriptome modules showed that there may be a modular structure to these effects, with specific pathogen SNPs linking to specific modules in either the host or the pathogen. However, future validation work will be required to further understand the directionality and mechanism of this crosstalk. For pathogen eQTL affecting host networks, mutants in the eQTL and the host target genes could elucidate whether the pathogen is specifically targeting host networks, or whether the host is sensing and countering the pathogen attack in response to particular signals. Similar work in other systems will help to build our functional knowledge of cross-kingdom communication between host and pathogen.

## METHODS

### Experimental design

We used a previously described collection of *B. cinerea* genotypes that were isolated as single spores from natural infections of fruit and vegetable tissues collected in California and internationally (Atwell, Corwin et al. 2015, Zhang, Corwin et al. 2017, Fordyce, Soltis et al. 2018, Zhang, Corwin et al. 2018). We focused analysis on the *A. thaliana* accession Columbia-0 (Col-0), and all plants were grown as described in a previous study, with 4-fold replication of the full randomized complete block experimental design across two independent experiments (Zhang, Corwin et al. 2017, Zhang, Corwin et al. 2018). The original study included wildtype Col-0 *A. thaliana* hosts, as well as knockouts to the salicylic acid pathway (*npr1-1*) and to jasmonic acid sensitivity (*coi1-1*). Leaves were harvested 5 weeks after sowing, and inoculated in a detached leaf assay with spores of each of 96 *B. cinerea* isolates (Zhang, Corwin et al. 2017, Zhang, Corwin et al. 2018). Whole leaves were sampled at 16 hours post inoculation, prior to visible lesion formation, and flash-frozen for RNA isolation.

### Expression analysis

RNASeq libraries were prepared as previously described (Kumar, Ichihashi et al. 2012, Zhang, Corwin et al. 2017, Zhang, Corwin et al. 2018). Briefly, we prepared mRNA from leaves frozen at 16 hours post inoculation, and pooled amplified, size-selected libraries into four replicate groups of 96 barcoded libraries. Sequencing was completed on a single Illumina HiSeq 2500 (San Diego, CA) lane as single 50bp reads at the U.C. Davis Genome Center-DNA Technologies Core (Davis, CA). Individual libraries were then separated by adapter index from fastq files, evaluated for read quality and overrepresentation (FastQC Version 0.11.3, www.bioinformatics.babraham.ac.uk/projects/), and trimmed (fastx, http://hannonlab.cshl.edu/fastx_toolkit/commandline.html). Reads were aligned to the *A. thaliana* TAIR10.25 cDNA reference genome, followed by the *B. cinerea* B05.10 cDNA reference genome, and we pulled gene counts (Langmead, Trapnell et al. 2009, Li, Handsaker et al. 2009, Van Kan, Stassen et al. 2017, Zhang, Corwin et al. 2017, Zhang, Corwin et al. 2018). We summed counts across gene models, and normalized gene counts as previously described (Zhang, Corwin et al. 2017, Zhang, Corwin et al. 2018).

We used as input the model-adjusted means per transcript from negative binomial linked generalized linear models in previously published studies in the *A. thaliana* transcriptome and *B. cinerea* transcriptome (Zhang, Corwin et al. 2017, Zhang, Corwin et al. 2018). *A. thaliana* and *B. cinerea* transcript phenotypes were from least square means of normalized gene counts in a negative binomial generalized linear model (nbGLM) (Zhang, Corwin et al. 2017, Zhang, Corwin et al. 2018). We calculated linear models from the transcript data including the effects of isolate and host genotype. We z-scaled all transcript profiles prior to GWA.

### Genome wide association

For GEMMA mapping, we used 95 isolates with a total of 237,878 SNPs against the *B. cinerea* B05.10 genome (Atwell, Corwin et al. 2018). We used haploid binary SNP calls with MAF > 0.20 and <20% missingness. We ran GEMMA once per phenotype, across 9,267 *B cinerea* gene expression profiles and 23,947 *A. thaliana* gene expression profiles.

### Genome wide association of permuted phenotypes

To validate SNPs as significantly associated with transcript variation, we performed a comparative analysis of randomized phenotypes. Taking each transcriptional profile, we randomized the assignment of phenotypes across the 96-isolate collection. This analysis includes 9,267 randomized *B. cinerea* phenotypes and 23,947 randomized *A. thaliana* phenotypes, one from each measured expression profile. We repeated this randomization in a 5x permutation. We ran GEMMA on each of these permutations, and plotted SNP p-value vs. position (Figure 1). To threshold our individual expression profile GEMMA outputs, we considered p-values below the average 5% permutation threshold as significant; p < 1.96e-05 for *B. cinerea* and p < 2.90e-05 for *A. thaliana*. Permutation approaches are often more effective than p-value thresholding for determining significance across GWA studies with many phenotypes (Evans and Cardon 2006).

### Defining significant hotspots

We plotted the number of transcripts linked to each SNP, summed across all 5 permutations, to calculate permuted hotspot size. For any SNPs that linked to permuted hotspots of over 5 transcripts in *B. cinerea* or 10 transcripts in *A. thaliana*, we removed these SNPs from downstream analysis as likely false positives. The maximum hotspot size across any of the 5 permutations was 11 genes in *B. cinerea* and 80 genes in *A. thaliana*. We then conservatively defined significant hotspots as SNP peaks exceeding 20 transcripts in *B. cinerea* and 150 transcripts in *A. thaliana*. We further annotated hotspot SNPs to the nearest gene within a 2kb window. The average LD decay in the B. cinerea genome is < 1kb, so we can be relatively confident of SNPs tagging particular genes at the hotspot peaks (Atwell, Corwin et al. 2018). Three genes are annotated to pairs of neighboring hotspots, the rest are unique genes. Two genes on chromosome 12 denoting hotspots from *A. thaliana* gene expression appear closely linked; in fact, they are separated by ~80kb on the *B. cinerea* genome.

### Annotation of gene ontology and network membership

*A. thaliana* co-expression analysis identified 131 genes across four major networks (Zhang, Corwin et al. 2017). Network architecture varied by plant host, but a constitutive core was conserved across *A. thaliana* genotypes. We compared our eQTL hotspots (both the gene at eQTL hotspot SNP and all associated transcript profiles) to the largest *A. thaliana* network lists (*npr1-1* background) to estimate all possible regulatory ties. We identified gene overlap with two of the major networks; Network I, camalexin biosynthesis; Network IV, chloroplast function.

*B. cinerea* co-expression analysis identified ten major co-expression networks containing 5 to 242 genes (Zhang, Corwin et al. 2018). We identified gene overlap with four of these networks, including one likely involved in fungal vesicle virulence processes including growth and toxin secretion (vesicle/ virulence), one involved in translation and protein synthesis (translation/ growth). These networks maintained a consistent core across the 3 *A. thaliana* host genotypes, but linkages varied; as such we compared our gene lists with the networks across all 12 hosts and included both host-dependent and host-independent annotations of our hotspots.

We annotated functions to B. cinerea genes using the BotPortal resource (http://dx.doi.org/10.15454/IHYJCX) and looked for patterns indicating signal peptides for secretion using the SignalP-5.0 Server (http://www.cbs.dtu.dk/services/SignalP-5.0/). We looked for functional overrepresentation among the genes targeted by each *A. thaliana* eQTL hotspot using the PANTHER overrepresentation test implemented by plant GO term enrichment from TAIR (Lamesch, Berardini et al. 2011, Mi, Muruganujan et al. 2013).

### Pathway focus

We focused further *cis*-effects analysis on three networks which were highly conserved across *B. cinerea* isolates (Zhang, Corwin et al. 2018). We clustered isolates by SNP data within focal networks. Hierarchical clustering was computed using the R package pvclust based on mean linkage (UPGMA), with correlation distance and 1000 bootstrap replications (Suzuki and Shimodaira 2015). AU p-values are reported in red, BP values in green. Edges with high AU values are considered strongly supported by the data, and clustering is drawn according to these edges with AU > 95%.

For botcinic acid biosynthesis, the major deletion extends 53.5 kb and includes SNP 4kb from the 5’ end of the chromosome, indicating a telomeric loss on chromosome 1. We selected a focal region encompassing the deletion endpoints (1.4029, 1.82614) and an additional 2 genes beyond the deletion boundaries (Bcin01g00170, Bcin01g00190) (Figure 3c). We removed 10 SNPs that were likely miscalled (SNP state ~ inverse compared to surrounding region) and called all SNPs within the deletion region as missing.

## Supporting information

Supplemental Data Set 2

Supplemental Data Set 3

Supplemental Data Set 4

## SUPPLEMENTAL FIGURE AND TABLE LEGENDS

**Supplemental Table 1. Annotation of the *B. cinerea* genetic targets of *B. cinerea* hotspots**. Columns include the hotspot SNP and the nearest gene, and all other columns pertain to the target gene modulated by the eQTL. Additional information about the target gene includes gene name, gene function, annotation as an enzyme, and target transcript. Gene functions are IPR numbers from the InterPro database. **Supplemental Data Set 1. Functional summary of the *B. cinerea* genetic targets of *B. cinerea* hotspots**. These count occurrences of major functional categories among the hotspot target genes.

**Supplemental Data Set 2. Annotation of the *A. thaliana* genetic targets of *B. cinerea* hotspots**. Columns include the *B. cinerea* hotspot SNP and the nearest *B. cinerea* gene, and the *A. thaliana* target gene.

**Supplemental Data Set 3. Gene ontology analysis of the *A. thaliana* genetic targets of *B. cinerea* hotspots**. These include all PANTHER overrepresentation test outputs for target gene sets within each eQTL hotspot. Hotspots are labeled by SNP and nearest *B. cinerea* gene. All calculations are from Bonferroni-corrected Fisher’s exact tests, and only significant GO categories are presented.

**Supplemental Data Set 4. Distance from B. cinerea transcripts to top SNP hits**. Dataset includes the lowest 50 p-values of SNP-transcript associations and the top 50 effect size estimates (beta) of SNP-transcript associations. Distance column indicates whether SNP and transcript are on separate chromosomes (*trans*), or if on the same chromosome (*cis*), the distance between the SNP and the transcript center.

**Figure S1. Distribution of number of associations and p-values for SNP-transcript associations**. *B. cinerea* transcripts with significant SNP associations (1,616 total) are a and c, *A. thaliana* transcripts with significant SNP associations (5,213 total) are b and d. In a and b, histograms depict the distribution of number of significant SNP associations per each transcript. In c and d, boxplots encompass the top 1 SNP associated with each transcript. Box edges delimit the first and third quartile, the thick center line delimits the median. Whiskers extend to 1.5 times the interquartile range and additional points indicate outliers.

**Figure S2. Manhattan-type plot of GEMMA results of transcriptome-wide *B. cinerea* expression phenotypes**. Each point represents a single transcript-SNP p-value of association. Panel a is a Manhattan-type plot of the top 1 SNP hit per *B. cinerea* transcript on Col-0 *A. thaliana*. Panel b is a Manhattan-type plot of the top 1 SNP hit per *A. thaliana* transcript when infected by *B. cinerea*.

**Figure S3. Distance between transcript center and top SNP location for all *B. cinerea* expression profiles on Col-0 *A. thaliana***. Distances are in Mb, including only top SNPs on the same chromosome as the focal gene. Panel a data include the top 1 SNP identified by GEMMA association with each transcript expression profile (lowest p-value for association). Panel b describes the length of individual chromosomes. Panel c data include the shortest distance between transcript genomic location and top 1 SNP identified by GEMMA association with each transcript expression profile (lowest p-value for association) out of 5 permutations.

**Figure S4. *cis*-effect analysis of the botrydial biosynthetic gene network**. Panel a is hierarchical clustering of *B. cinerea* isolates from SNPs within the botrydial biosynthetic gene network. Clustering was based on mean linkage (UPGMA), with correlation distance and 1000 bootstrap replications. AU p-values are reported in red, BP values in green. Edges with high AU values are considered strongly supported by the data, and clustering is drawn according to these edges with AU > 95%. Panel b is Violin plots of botrydial network-level expression within *B. cinerea* clusters. Isolates are clustered based membership in groups defined by hierarchical clustering of the SNPs within the botrydial biosynthesis network.

**Figure S5. *cis*-effect analysis of the cyclic peptide biosynthetic gene network**. Panel a is hierarchical clustering of *B. cinerea* isolates from SNPs within the cyclic peptide biosynthetic gene network. Clustering was based on mean linkage (UPGMA), with correlation distance and 1000 bootstrap replications. AU p-values are reported in red, BP values in green. Edges with high AU values are considered strongly supported by the data, and clustering is drawn according to these edges with AU > 95%. Panel b is Violin plots of cyclic peptide network-level expression within *B. cinerea* clusters. Isolates are clustered based membership in groups defined by hierarchical clustering of the SNPs within the botrydial biosynthesis network.

**Figure S6. Interspecific hotspot comparison on the *B. cinerea* genome with the top 10 genes per SNP**. For each SNP that is a top hit for one or more transcripts, the number of associated transcripts is counted, across both the *B. cinerea* transcriptome and the *A. thaliana* transcriptome.

## REFERENCES

Allen, M., M. M. Carrasquillo, C. Funk, B. D. Heavner, F. Zou, C. S. Younkin, J. D. Burgess, H.-S. Chai, J. Crook and J. A. Eddy (2016). “Human whole genome genotype and transcriptome data for Alzheimer’s and other neurodegenerative diseases.” Scientific data 3: 160089.

Atwell, S., J. Corwin, N. Soltis and D. Kliebenstein (2018). “Resequencing and association mapping of the generalist pathogen Botrytis cinerea.” bioRxiv.

Atwell, S., J. Corwin, N. Soltis, A. Subedy, K. Denby and D. J. Kliebenstein (2015). “Whole genome resequencing of Botrytis cinerea isolates identifies high levels of standing diversity.” Frontiers in microbiology 6: 996.

Barrett, L. G., J. M. Kniskern, N. Bodenhausen, W. Zhang and J. Bergelson (2009). “Continua of specificity and virulence in plant host–pathogen interactions: causes and consequences.” New Phytologist 183(3): 513–529.

Bartha, I., P. J. McLaren, C. Brumme, R. Harrigan, A. Telenti and J. Fellay (2017). “Estimating the respective contributions of human and viral genetic variation to HIV control.” PLoS computational biology 13(2): e1005339.

Bartoli, C. and F. Roux (2017). “Genome-Wide Association Studies In Plant Pathosystems: Toward an Ecological Genomics Approach.” Frontiers in plant science 8.

Bilgin, D. D., J. A. Zavala, J. Zhu, S. J. Clough, D. R. Ort and E. H. DeLUCIA (2010). “Biotic stress globally downregulates photosynthesis genes.” Plant, cell & environment 33(10): 1597–1613.

Brem, R. B., G. Yvert, R. Clinton and L. Kruglyak (2002). “Genetic dissection of transcriptional regulation in budding yeast.” Science 296(5568): 752–755.

Caseys, C., G. Shi, N. Soltis, R. Gwinner, J. Corwin, S. Atwell and D. Kliebenstein (2018). “A generalist pathogen view of plant evolution.” bioRxiv: 507491.

Chan, E. K., H. C. Rowe, B. G. Hansen and D. J. Kliebenstein (2010). “The complex genetic architecture of the metabolome.” PLoS Genet 6(11): e1001198.

Chen, X., C. A. Hackett, R. E. Niks, P. E. Hedley, C. Booth, A. Druka, T. C. Marcel, A. Vels, M. Bayer and I. Milne (2010). “An eQTL analysis of partial resistance to Puccinia hordei in barley.” PLoS One 5(1): e8598.

Christie, N., A. A. Myburg, F. Joubert, S. L. Murray, M. Carstens, Y. C. Lin, J. Meyer, B. G. Crampton, S. A. Christensen and J. F. Ntuli (2017). “Systems genetics reveals a transcriptional network associated with susceptibility in the maize–grey leaf spot pathosystem.” The Plant Journal 89(4): 746–763.

Colmenares, A. J., J. Aleu, R. Duran-Patron, I. G. Collado and R. Hernandez-Galan (2002). “The putative role of botrydial and related metabolites in the infection mechanism of Botrytis cinerea.” Journal of chemical ecology 28(5): 997–1005.

Corwin, J. A., D. Copeland, J. Feusier, A. Subedy, R. Eshbaugh, C. Palmer, J. Maloof and D. J. Kliebenstein (2016). “The quantitative basis of the Arabidopsis innate immune system to endemic pathogens depends on pathogen genetics.” PLoS Genet 12(2): e1005789.

Corwin, J. A., A. Subedy, R. Eshbaugh and D. J. Kliebenstein (2016). “Expansive phenotypic landscape of Botrytis cinerea shows differential contribution of genetic diversity and plasticity.” Molecular Plant-Microbe Interactions 29(4): 287–298.

Cui, H., K. Tsuda and J. E. Parker (2015). “Effector-triggered immunity: from pathogen perception to robust defense.” Annual review of plant biology 66: 487–511.

Deighton, N., I. Muckenschnabel, A. J. Colmenares, I. G. Collado and B. Williamson (2001). “Botrydial is produced in plant tissues infected by Botrytis cinerea.” Phytochemistry 57(5): 689–692.

Denby, K. J., P. Kumar and D. J. Kliebenstein (2004). “Identification of Botrytis cinerea susceptibility loci in Arabidopsis thaliana.” The Plant Journal 38(3): 473–486.

Dong, S., S. Raffaele and S. Kamoun (2015). “The two-speed genomes of filamentous pathogens: waltz with plants.” Current opinion in genetics & development 35: 57–65.

Evans, D. M. and L. R. Cardon (2006). “Genome-wide association: a promising start to a long race.” Trends in Genetics 22(7): 350–354.

Fordyce, R., N. Soltis, C. Caseys, G. Gwinner, J. Corwin, S. Atwell, D. Copeland, J. Feusier, A. Subedy, R. Eshbaugh and D. Kliebenstein (2018). “Combining Digital Imaging and GWA Mapping to Dissect Visual Traits in Plant/Pathogen Interactions.” Plant Physiology.

Giraldo, M. C. and B. Valent (2013). “Filamentous plant pathogen effectors in action.” Nature Reviews Microbiology 11(11): 800.

Glazebrook, J. (2005). “Contrasting mechanisms of defense against biotrophic and necrotrophic pathogens.” Annu. Rev. Phytopathol. 43: 205–227.

Goss, E. M. and J. Bergelson (2006). “Variation in resistance and virulence in the interaction between Arabidopsis thaliana and a bacterial pathogen.” Evolution 60(8): 1562–1573.

Guo, Y., S. Fudali, J. Gimeno, P. DiGennaro, S. Chang, V. M. Williamson, D. M. Bird and D. M. Nielsen (2017). “Networks underpinning symbiosis revealed through cross-species eQTL mapping.” Genetics: genetics. 117.202531.

Hsu, J. and J. D. Smith (2012). “Genome wide studies of gene expression relevant to coronary artery disease.” Current opinion in cardiology 27(3): 210.

Jiang, Z., F. He and Z. Zhang (2017). “Large-scale transcriptome analysis reveals arabidopsis metabolic pathways are frequently influenced by different pathogens.” Plant molecular biology 94(4–5): 453–467.

Keurentjes, J. J., J. Fu, I. R. Terpstra, J. M. Garcia, G. van den Ackerveken, L. B. Snoek, A. J. Peeters, D. Vreugdenhil, M. Koornneef and R. C. Jansen (2007). “Regulatory network construction in Arabidopsis by using genome-wide gene expression quantitative trait loci.” Proceedings of the National Academy of Sciences 104(5): 1708–1713.

Kou, Y. and S. Wang (2010). “Broad-spectrum and durability: understanding of quantitative disease resistance.” Current opinion in plant biology 13(2): 181–185.

Kumar, R., Y. Ichihashi, S. Kimura, D. H. Chitwood, L. R. Headland, J. Peng, J. N. Maloof and N. R. Sinha (2012). “A high-throughput method for Illumina RNA-Seq library preparation.” Frontiers in plant science 3.

Lamesch, P., T. Z. Berardini, D. Li, D. Swarbreck, C. Wilks, R. Sasidharan, R. Muller, K. Dreher, D. L. Alexander and M. Garcia-Hernandez (2011). “The Arabidopsis Information Resource (TAIR): improved gene annotation and new tools.” Nucleic acids research 40(D1): D1202–D1210.

Langmead, B., C. Trapnell, M. Pop and S. L. Salzberg (2009). “Ultrafast and memory-efficient alignment of short DNA sequences to the human genome.” Genome biology 10(3): R25.

Lannou, C. (2012). “Variation and selection of quantitative traits in plant pathogens.” Annual Review of Phytopathology 50: 319–338.

Li, H., B. Handsaker, A. Wysoker, T. Fennell, J. Ruan, N. Homer, G. Marth, G. Abecasis and R. Durbin (2009). “The sequence alignment/map format and SAMtools.” Bioinformatics 25(16): 2078–2079.

Lo Presti, L., D. Lanver, G. Schweizer, S. Tanaka, L. Liang, M. Tollot, A. Zuccaro, S. Reissmann and R. Kahmann (2015). “Fungal effectors and plant susceptibility.” Annual review of plant biology 66: 513–545.

Marone, D., M. Russo, G. Laidò, A. De Leonardis and A. Mastrangelo (2013). “Plant nucleotide binding site–leucine-rich repeat (NBS-LRR) genes: active guardians in host defense responses.” International journal of molecular sciences 14(4): 7302–7326.

Martínez-Soto, D., A. M. Robledo-Briones, A. A. Estrada-Luna and J. Ruiz-Herrera (2013). “Transcriptomic analysis of U stilago maydis infecting Arabidopsis reveals important aspects of the fungus pathogenic mechanisms.” Plant signaling & behavior 8(8): e25059.

Meng, X. and S. Zhang (2013). “MAPK cascades in plant disease resistance signaling.” Annual review of phytopathology 51: 245–266.

Mi, H., A. Muruganujan, J. T. Casagrande and P. D. Thomas (2013). “Large-scale gene function analysis with the PANTHER classification system.” Nature protocols 8(8): 1551.

Monks, S., A. Leonardson, H. Zhu, P. Cundiff, P. Pietrusiak, S. Edwards, J. Phillips, A. Sachs and E. Schadt (2004). “Genetic inheritance of gene expression in human cell lines.” The American Journal of Human Genetics 75(6): 1094–1105.

Nobori, T., A. C. Velásquez, J. Wu, B. H. Kvitko, J. M. Kremer, Y. Wang, S. Y. He and K. Tsuda (2018). “Transcriptome landscape of a bacterial pathogen under plant immunity.” Proceedings of the National Academy of Sciences 115(13): E3055–E3064.

Nomura, K., M. Melotto and S.-Y. He (2005). “Suppression of host defense in compatible plant– Pseudomonas syringae interactions.” Current opinion in plant biology 8(4): 361–368.

Pinedo, C., C.-M. Wang, J.-M. Pradier, B. Dalmais, M. Choquer, P. Le Pêcheur, G. Morgant, I. G. Collado, D. E. Cane and M. Viaud (2008). “Sesquiterpene synthase from the botrydial biosynthetic gene cluster of the phytopathogen Botrytis cinerea.” ACS chemical biology 3(12): 791–801.

Poland, J. A., P. J. Balint-Kurti, R. J. Wisser, R. C. Pratt and R. J. Nelson (2009). “Shades of gray: the world of quantitative disease resistance.” Trends in plant science 14(1): 21–29.

Porquier, A., J. Moraga, G. Morgant, B. Dalmais, A. Simon, H. Sghyer, I. G. Collado and M. Viaud (2019). “Botcinic acid biosynthesis in Botrytis cinerea relies on a subtelomeric gene cluster surrounded by relics of transposons and is regulated by the Zn 2 Cys 6 transcription factor BcBoa13.” Current genetics: 1–16.

Porquier, A., G. Morgant, J. Moraga, B. Dalmais, I. Luyten, A. Simon, J.-M. Pradier, J. Amselem, I. G. Collado and M. Viaud (2016). “The botrydial biosynthetic gene cluster of Botrytis cinerea displays a bipartite genomic structure and is positively regulated by the putative Zn (II) 2Cys6 transcription factor BcBot6.” Fungal genetics and biology 96: 33–46.

Rivas, M. A., M. Beaudoin, A. Gardet, C. Stevens, Y. Sharma, C. K. Zhang, G. Boucher, S. Ripke, D. Ellinghaus and N. Burtt (2011). “Deep resequencing of GWAS loci identifies independent rare variants associated with inflammatory bowel disease.” Nature genetics 43(11): 1066.

Roux, F., D. Voisin, T. Badet, C. Balagué, X. Barlet, C. Huard-Chauveau, D. Roby and S. Raffaele (2014). “Resistance to phytopathogens e tutti quanti: placing plant quantitative disease resistance on the map.” Molecular plant pathology 15(5): 427–432.

Rowe, H. C. and D. J. Kliebenstein (2008). “Complex genetics control natural variation in Arabidopsis thaliana resistance to Botrytis cinerea.” Genetics 180(4): 2237–2250.

Schadt, E. E., S. A. Monks, T. A. Drake, A. J. Lusis, N. Che, V. Colinayo, T. G. Ruff, S. B. Milligan, J. R. Lamb and G. Cavet (2003). “Genetics of gene expression surveyed in maize, mouse and man.” Nature 422(6929): 297.

Siewers, V., M. Viaud, D. Jimenez-Teja, I. G. Collado, C. S. Gronover, J.-M. Pradier, B. Tudzynsk and P. Tudzynski (2005). “Functional analysis of the cytochrome P450 monooxygenase gene bcbot1 of Botrytis cinerea indicates that botrydial is a strain-specific virulence factor.” Molecular plant-microbe interactions 18(6): 602–612.

Soltis, N. E., S. Atwell, G. Shi, R. F. Fordyce, R. Gwinner, D. Gao, A. Shafi and D. J. Kliebenstein (2019). “Interactions of tomato and Botrytis genetic diversity: Parsing the contributions of host differentiation, domestication and pathogen variation.” The Plant Cell: tpc.00857.02018.

St. Clair, D. A. (2010). “Quantitative disease resistance and quantitative resistance loci in breeding.” Annual review of phytopathology 48: 247–268.

Suzuki, R. and H. Shimodaira (2015). “pvclust: Hierarchical Clustering with P-Values via Multiscale Bootstrap Resampling..” R package version 2.0-0.

Valero-Jiménez, C. A., J. Veloso, M. Staats and J. A. van Kan (2019). “Comparative genomics of plant pathogenic Botrytis species with distinct host specificity.” BMC Genomics 20(1): 203.

Van Kan, J. A., J. H. Stassen, A. Mosbach, T. A. Van Der Lee, L. Faino, A. D. Farmer, D. G. Papasotiriou, S. Zhou, M. F. Seidl and E. Cottam (2017). “A gapless genome sequence of the fungus Botrytis cinerea.” Molecular plant pathology 18(1): 75–89.

Visscher, P. M., N. R. Wray, Q. Zhang, P. Sklar, M. I. McCarthy, M. A. Brown and J. Yang (2017). “10 years of GWAS discovery: biology, function, and translation.” The American Journal of Human Genetics 101(1): 5–22.

Wang, M., F. Roux, C. Bartoli, C. Huard-Chauveau, C. Meyer, H. Lee, D. Roby, M. S. McPeek and J. Bergelson (2018). “Two-way mixed-effects methods for joint association analysis using both host and pathogen genomes.” Proceedings of the National Academy of Sciences 115(24): E5440–E5449.

West, M. A. L., K. Kim, D. J. Kliebenstein, H. van Leeuwen, R. W. Michelmore, R. W. Doerge and D. A. St.Clair (2007). “Global eQTL mapping reveals the complex genetic architecture of transcript level variation in Arabidopsis.” Genetics 175: 1441–1450.

Wu, J., B. Cai, W. Sun, R. Huang, X. Liu, M. Lin, S. Pattaradilokrat, S. Martin, Y. Qi and S. C. Nair (2015). “Genome-wide analysis of host-Plasmodium yoelii interactions reveals regulators of the type I interferon response.” Cell reports 12(4): 661–672.

Wu, J. Q., S. Sakthikumar, C. Dong, P. Zhang, C. A. Cuomo and R. F. Park (2017). “Comparative genomics integrated with association analysis identifies candidate effector genes corresponding to Lr20 in phenotype-paired Puccinia triticina isolates from Australia.” Frontiers in plant science 8.

Zhang, W., J. A. Corwin, D. Copeland, J. Feusier, R. Eshbaugh, F. Chen, S. Atwell and D. J. Kliebenstein (2017). “Plastic transcriptomes stabilize immunity to pathogen diversity: the jasmonic acid and salicylic acid networks within the Arabidopsis/Botrytis pathosystem.” The Plant Cell: tpc. 00348.02017.

Zhang, W., J. A. Corwin, D. Copeland, J. Feusier, R. Eshbaugh, D. E. Cook, S. Atwell and D. J. Kliebenstein (2018). “Network connections across kingdoms illuminate a potential metabolic battlefield.” bioRxiv.

Zhou, X. and M. Stephens (2012). “Genome-wide efficient mixed-model analysis for association studies.” Nature genetics 44(7): 821.

Zou, F., H. S. Chai, C. S. Younkin, M. Allen, J. Crook, V. S. Pankratz, M. M. Carrasquillo, C. N. Rowley, A. A. Nair and S. Middha (2012). “Brain expression genome-wide association study (eGWAS) identifies human disease-associated variants.” PLoS genetics 8(6): e1002707.

