## Supplemental Figures S1 to S6 and Supplemental Table 1 for "Pathogen genetic control of transcriptome variation in the *Arabidopsis thaliana* – *Botrytis cinerea* pathosystem"

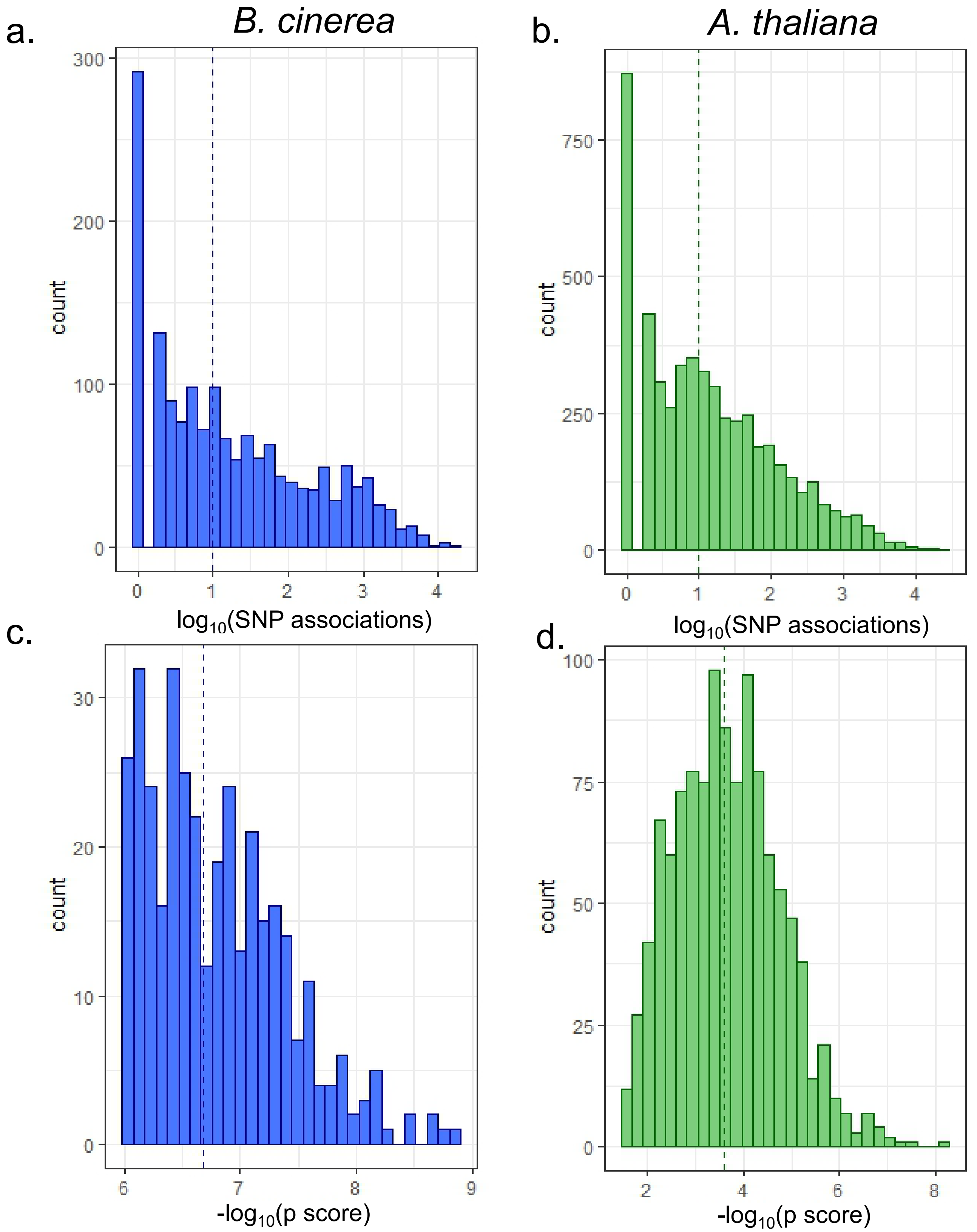


**Figure S1. Distribution of number of associations and p-values for SNP-transcript associations**. *B. cinerea* transcripts with significant SNP associations (1,616 total) are a and c, *A. thaliana* transcripts with significant SNP associations (5,213 total) are b and d. In a and b, histograms depict the distribution of number of significant SNP associations per each transcript. In c and d, boxplots encompass the top 1 SNP associated with each transcript. Box edges delimit the first and third quartile, the thick center line delimits the median. Whiskers extend to 1.5 times the interquartile range and additional points indicate outliers.

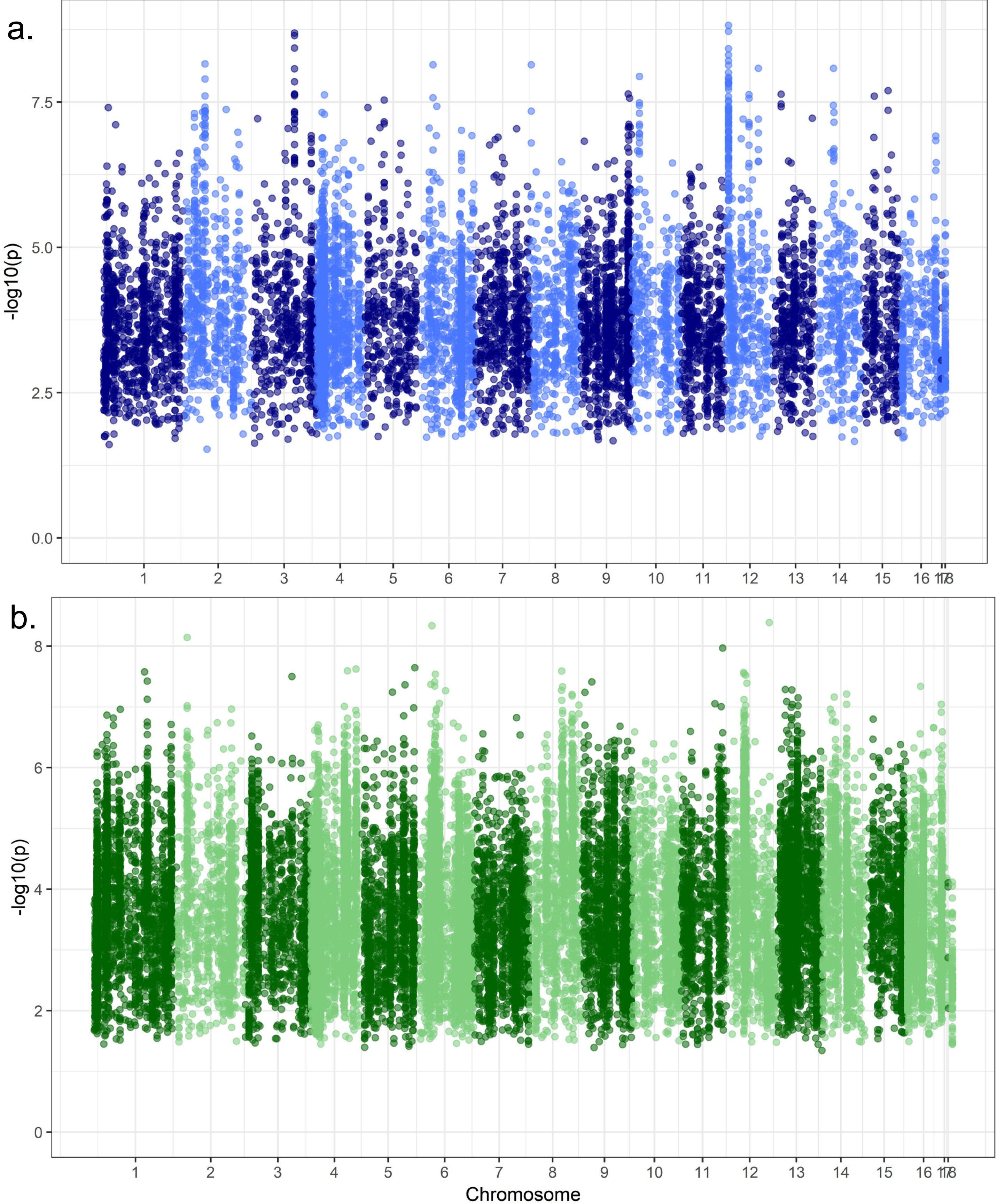


**Figure S2. Manhattan-type plot of GEMMA results of transcriptome-wide *B. cinerea* expression phenotypes.** Each point represents a single transcript-SNP p-value of association. Panel a is a Manhattan-type plot of the top 1 SNP hit per *B. cinerea* transcript on Col-0 *A. thaliana*. Panel b is a Manhattan-type plot of the top 1 SNP hit per *A. thaliana* transcript when infected by *B. cinerea*.

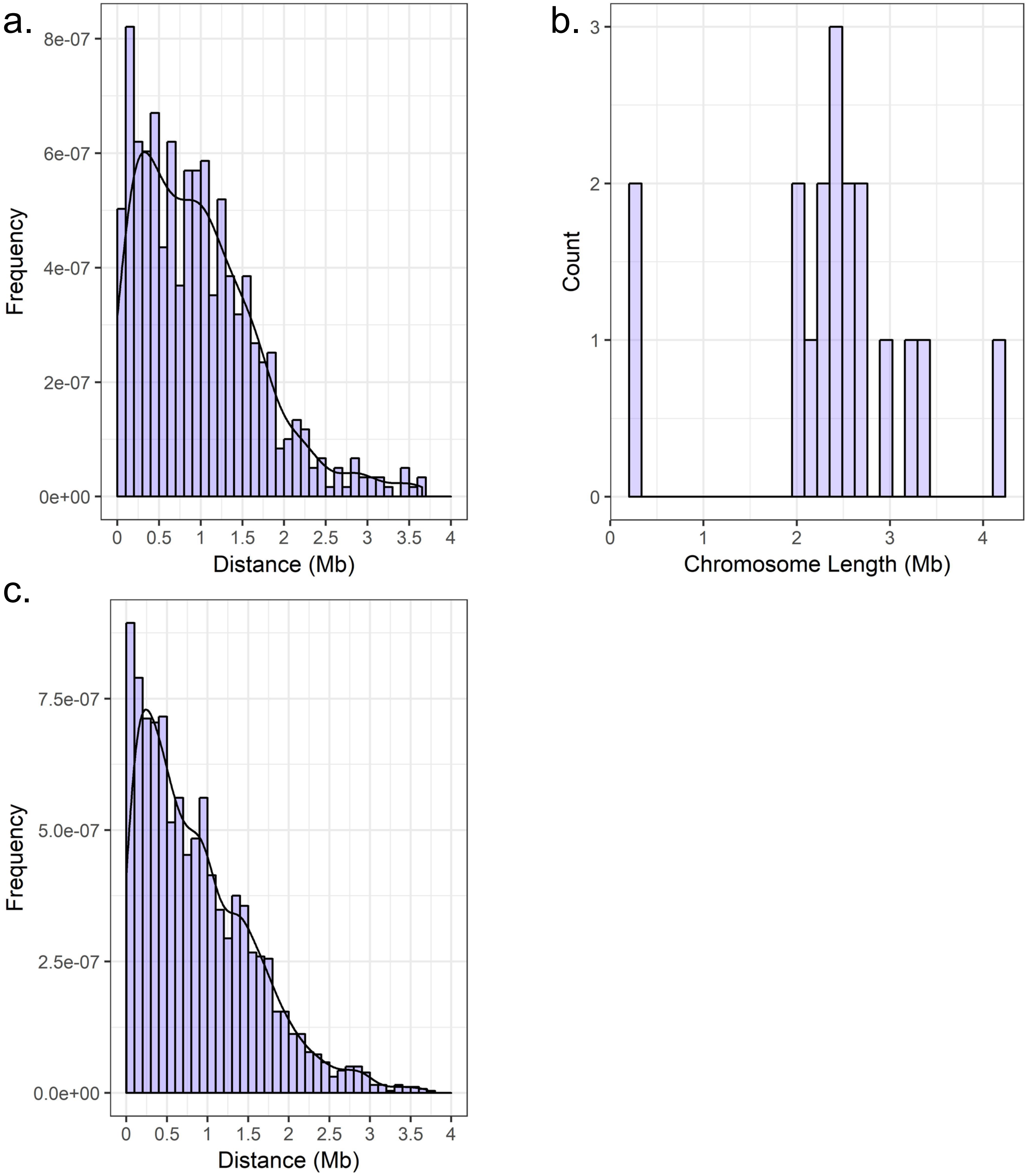


**Figure S3. Distance between transcript center and top SNP location for all *B. cinerea* expression profiles on Col-0 *A. thaliana*.** Distances are in Mb, including only top SNPs on the same chromosome as the focal gene. Panel a data include the top 1 SNP identified by GEMMA association with each transcript expression profile (lowest p-value for association). Panel b describes the length of individual chromosomes. Panel c data include the shortest distance between transcript genomic location and top 1 SNP identified by GEMMA association with each transcript expression profile (lowest p-value for association) out of 5 permutations.

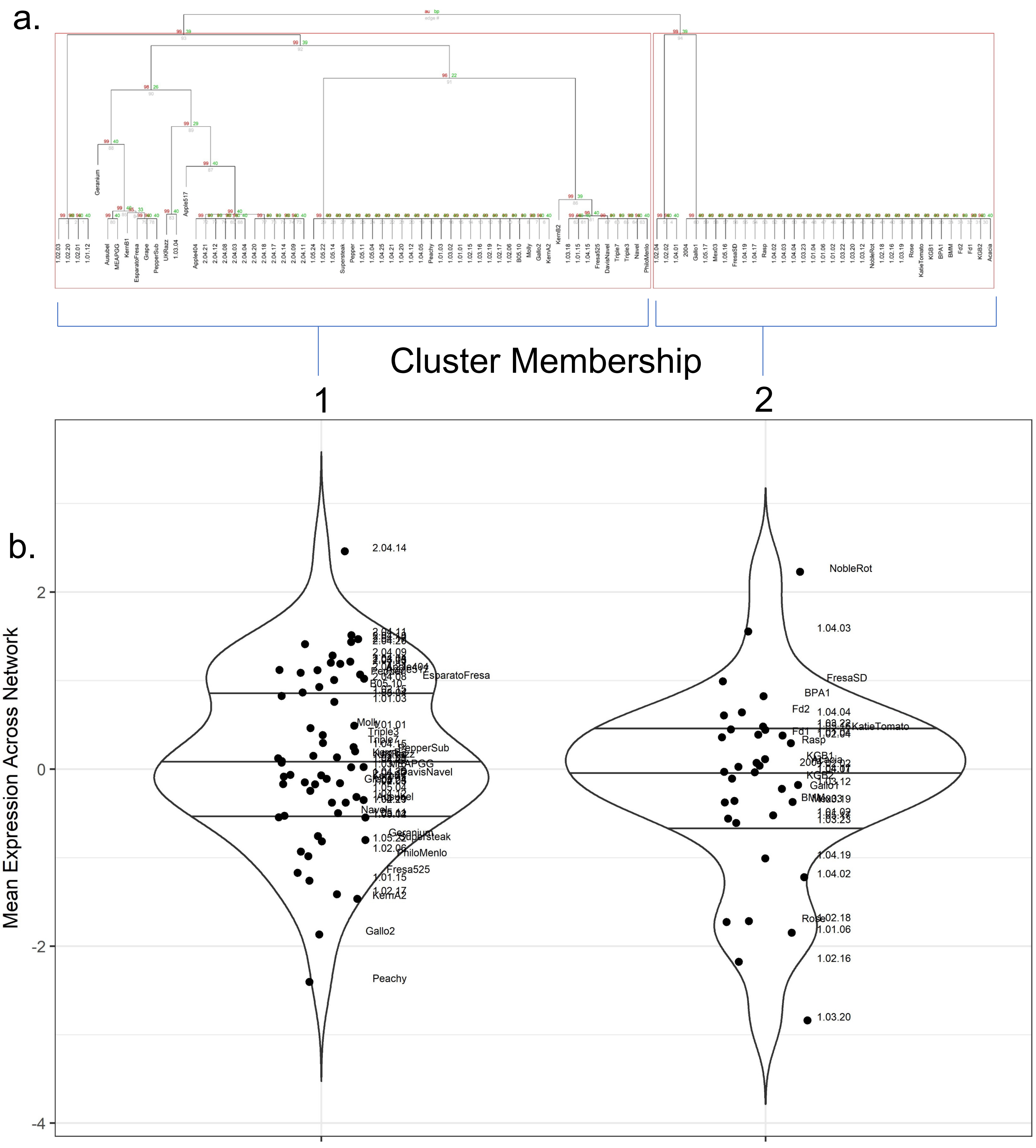


**Figure S4. *cis*-effect analysis of the botrydial biosynthetic gene network.** Panel a is hierarchical clustering of *B. cinerea* isolates from SNPs within the botrydial biosynthetic gene network. Clustering was based on mean linkage (UPGMA), with correlation distance and 1000 bootstrap replications. AU p-values are reported in red, BP values in green. Edges with high AU values are considered strongly supported by the data, and clustering is drawn according to these edges with AU > 95%. Panel b is Violin plots of botrydial network-level expression within *B. cinerea* clusters. Isolates are clustered based membership in groups defined by hierarchical clustering of the SNPs within the botrydial biosynthesis network.

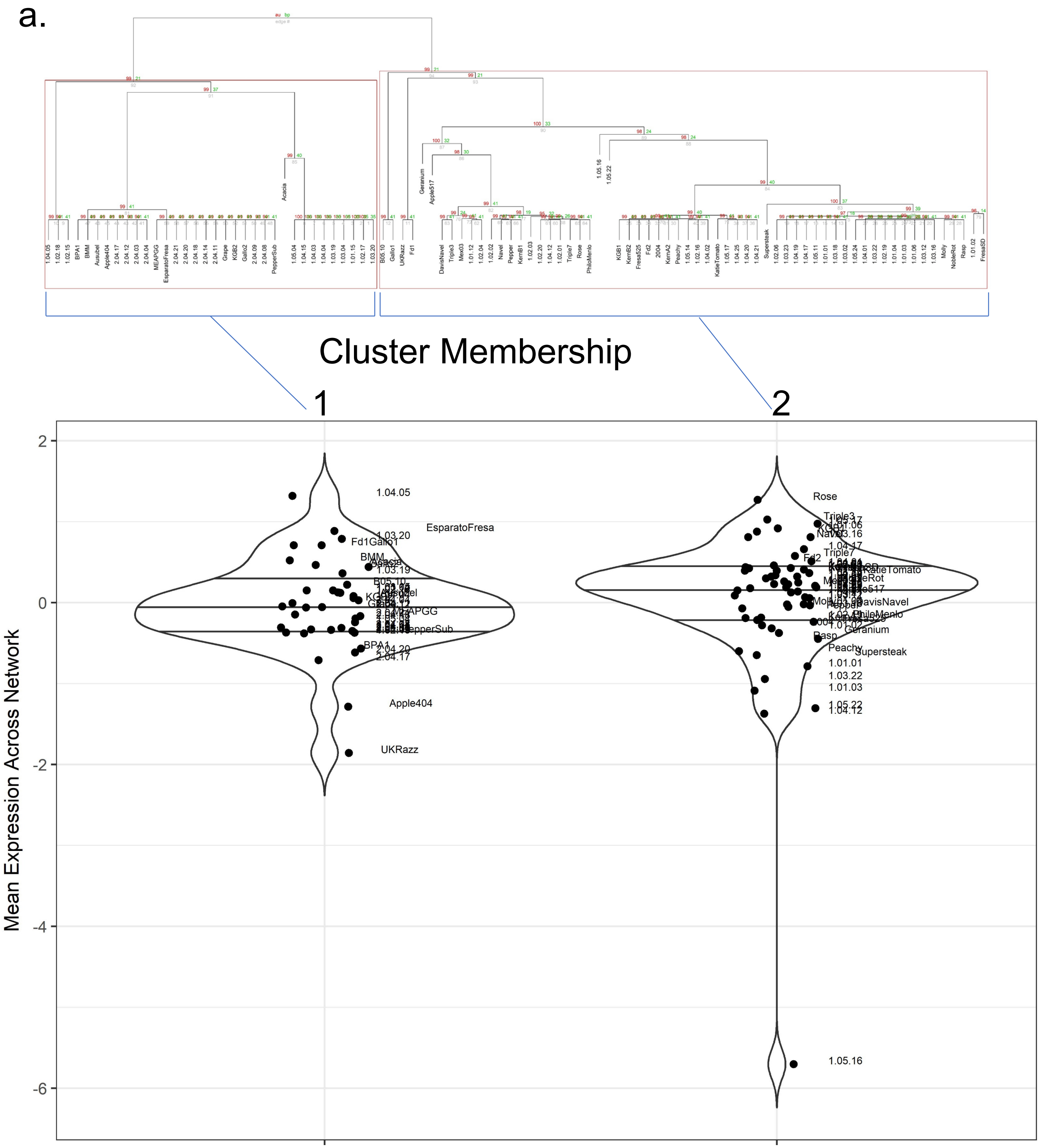


**Figure S5. *cis*-effect analysis of the cyclic peptide biosynthetic gene network.** Panel a is hierarchical clustering of *B. cinerea* isolates from SNPs within the cyclic peptide biosynthetic gene network. Clustering was based on mean linkage (UPGMA), with correlation distance and 1000 bootstrap replications. AU p-values are reported in red, BP values in green. Edges with high AU values are considered strongly supported by the data, and clustering is drawn according to these edges with AU > 95%. Panel b is Violin plots of cyclic peptide network-level expression within *B. cinerea* clusters. Isolates are clustered based membership in groups defined by hierarchical clustering of the SNPs within the botrydial biosynthesis network.

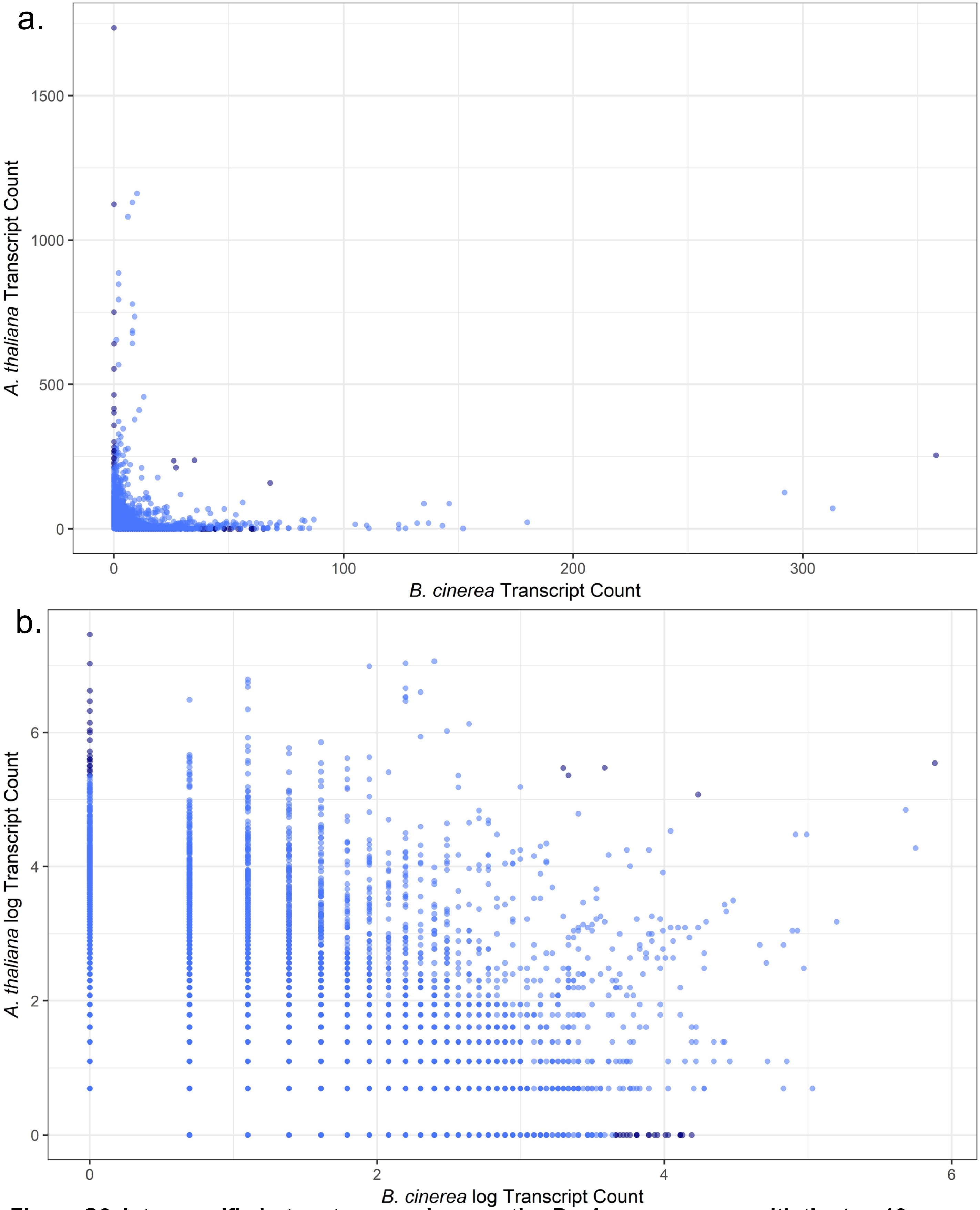


**Figure S6. Interspecific hotspot comparison on the *B. cinerea* genome with the top 10 genes per SNP.** For each SNP that is a top hit for one or more transcripts, the number of associated transcripts is counted, across both the *B. cinerea* transcriptome and the *A. thaliana* transcriptome.

**Supplemental Table 1. Functional summary of the *B. cinerea* genetic targets of *B. cinerea* hotspots.** These count occurrences of major functional categories among the hotspot target genes.

| | |
|---|---|
| **Total annotated** | **412** |
| **Enzyme** | 140 |
| **Heterokaryon domain** | 1 |
| **Gene with a peak of expression during the outset of colonization** | 2 |
| **TF** | 11 |
| **Gene induced in early infection stage** | 1 |
| **Expressed in planta** | 1 |
| **Mitogen-activated protein kinase** | 2 |
